# Snail maintains the stem/progenitor state of skin epithelial cells and carcinomas through the autocrine effect of the matricellular protein Mindin

**DOI:** 10.1101/2021.06.26.450022

**Authors:** Krithika Badarinath, Binita Dam, Sunny Kataria, Ravindra K. Zirmire, Rakesh Dey, Randhir Singh, Tafheem A. Masudi, Janani Sambath, Prashanth Kumar, Akash Gulyani, You-Wen He, Sudhir Krishna, Colin Jamora

## Abstract

Intratumoral heterogeneity poses a major challenge in designing effective anti-cancer strategies. Accumulating evidence suggests that this heterogeneity arises from cancer stem cells (CSCs) that also drives tumor aggressiveness and drug resistance. The stemness of CSCs are preserved by an ill-defined combination of intrinsic and external factors and is particularly intriguing since they exist within a sea of similar cells at various degrees of differentiation. In models of cutaneous squamous cell carcinoma (cSCC), we discovered a non-EMT function for the transcription factor Snail in maintaining stemness of keratinocytes. This is accomplished by the secretion of the matricellular protein Mindin from Snail expressing cells, which creates a protective niche that impedes differentiation. In an autocrine fashion, extracellular Mindin activates a Src –STAT3 pathway to reinforce the stem/progenitor phenotype and disruption of this signalling module in human cSCC attenuates tumorigenesis. The expression of Mindin in multiple carcinomas, and its critical role in cancer progression suggests that it would be a promising target for therapeutic intervention.

## Introduction

Intratumoral heterogeneity, is a term to describe the diversification of both the non-malignant and malignant compartments of the tumor. The former is a local microenvironment, also known as the stroma, that is composed of different cell populations including fibroblasts, endothelial cells, and immune cells. There is burgeoning interest in the role of the tumor stroma which has a profound impact on a wide spectrum of tumorigenic processes ranging from metastasis (Guo and Deng, 2018) to therapeutic resistance (Straussman et al., 2012). Another aspect of heterogeneity arises among the malignant epithelial cells that comprise the tumor mass. These epithelial cells differ in their growth rates, tumorigenic potential, and differentiation status and are driven by alterations in their DNA sequence, epigenome, transcriptome, proteome, and metabolome (Somasundaram et al., 2012). A plethora of studies suggest that intratumor heterogeneity is also a leading determinant of treatment failure and poor overall survival of cancer patients, and is thus the subject of intense research from a biological as well as a treatment point of view. (Michor and Polyak, 2010; Somasundaram et al., 2012; Tellez-Gabriel et al., 2016).

A prominent model to explain the manifestation of intratumor heterogeneity is the cancer stem cell (CSC) concept introduced in the late 1990’s (Lapidot et al., 1994). The CSC hypothesis posits that the tumor is a hierarchically organised tissue wherein these tumor initiating cells generate the various types of differentiated cells of the tumor mass. Similar to normal somatic stem cells, CSCs are distinguished by their capacity for indefinite self-renewal, which positions them as a central player in cancer initiation, maintenance, and progression. Interestingly, the CSC population is not static but increases in number with tumor grade thereby increasing tumor heterogeneity (Li et al., 2013; Yu, 2012). Moreover, CSCs are usually quiescent, have increased proficiency in DNA repair, disable their apoptotic pathways, and express increased levels of efflux transporters (Phi et al., 2018). These characteristics render CSCs capable of evading standard cytotoxic therapies (Li et al., 2021). Another similarity with tissue stem cells is the residence of CSCs within a niche. This niche is a specialized microenvironment that regulates stem cell behaviour by providing cues in the form of both cellular contacts and soluble factors (Sun et al., 2019). These soluble factors can arise from the tumor cell itself as well as from cells in the tumor stroma (Plaks et al., 2015). Examples of growth factors and cytokines that maintain the undifferentiated state of cells are Hedgehog, Wnt, and Notch, while stemness-promoting intracellular signalling pathways utilize JAK/STAT, PI3K/phosphatase, Hippo and NF-κB (Matsui, 2016; Plaks et al., 2015).

Given the importance of CSCs, many studies have focused on understanding their role and regulation in tumorigenesis. One open question is how are the few CSCs maintained in a tumor mass amongst a sea of differentiating cells. To answer this question, we utilized a transgenic mouse model based on the overexpression of Snail in epidermal keratinocytes (*K14-Snail Tg*), which reproduces many of the cardinal features of squamous cell carcinoma (Du et al., 2010). Snail, is a transcription factor that is well known for its function in inducing an epithelial-mesenchymal transition (EMT) during embryogenesis (Murray and Gridley, 2006) and hair morphogenesis (Jamora et al., 2005), and is overexpressed in many carcinomas (Fan et al., 2012a; Goossens et al., 2017; Mani et al., 2009). Interestingly, there is a strong correlation between Snail expression with the number of CSCs (Hojo et al., 2018a; Ma et al., 2017; Zhou et al., 2014) as well as tumor aggressiveness (Smith et al., 2014; Zheng et al., 2015). Several proposals have been formulated to explain the connection between Snail and CSCs including the ability of EMTs to promote de-differentiation of cells to acquire stem cell properties (Wang and Unternaehrer, 2019). Moreover, extracellular signals such as TGFβ and Wnt, that are known to promote CSC production, are dependent on EMT drivers such as Snail and Twist (Scheel et al., 2011). Despite these correlations, the molecular mechanism by which Snail specifically regulates the stemness and number of CSCs remains largely unknown.

## Results

### Overexpression of *Snail* reinforces the stem/progenitor characteristics in epidermal keratinocytes *in vitro*

Given Snail’s canonical function in inducing an EMT, we tested if this process was activated in the *K14-Snail* transgenic (Snail Tg) skin by three different criteria: 1) lineage tracing assay (Tan et al., manuscript in preparation), 2) immunofluorescence to examine cells co-expressing markers of the epithelial (keratin 5) and mesenchymal cells (vimentin [Figure 1A] and collagen [Figure S1A]) and 3) maintenance of distinct epithelial and mesenchymal compartments delineated by the basement membrane marked by laminin 5 (Figure 1A). Altogether, this analysis unexpectedly revealed that transgenic expression of *Snail* in normal epidermal keratinocytes *in vivo* is insufficient to induce an EMT. To validate these surprising observations, we examined whether primary keratinocytes isolated from the Snail Tg skin exhibited features of EMT in vitro. We computed the EMT score for each sample using transcriptome data to estimate the EMT phenotype (Tan et al., 2014) (Figure 1B). The EMT scores of both wild type and Snail Tg cells were distributed between −0.18 to −0.20 indicating that the EMT status did not differ between the samples (Figure S1B), and both cell populations are epithelial in nature. Consistent with the *in vivo* data, primary keratinocytes isolated from the Snail Tg skin retain their epithelial characteristics *in vitro*.

**Figure 1:**
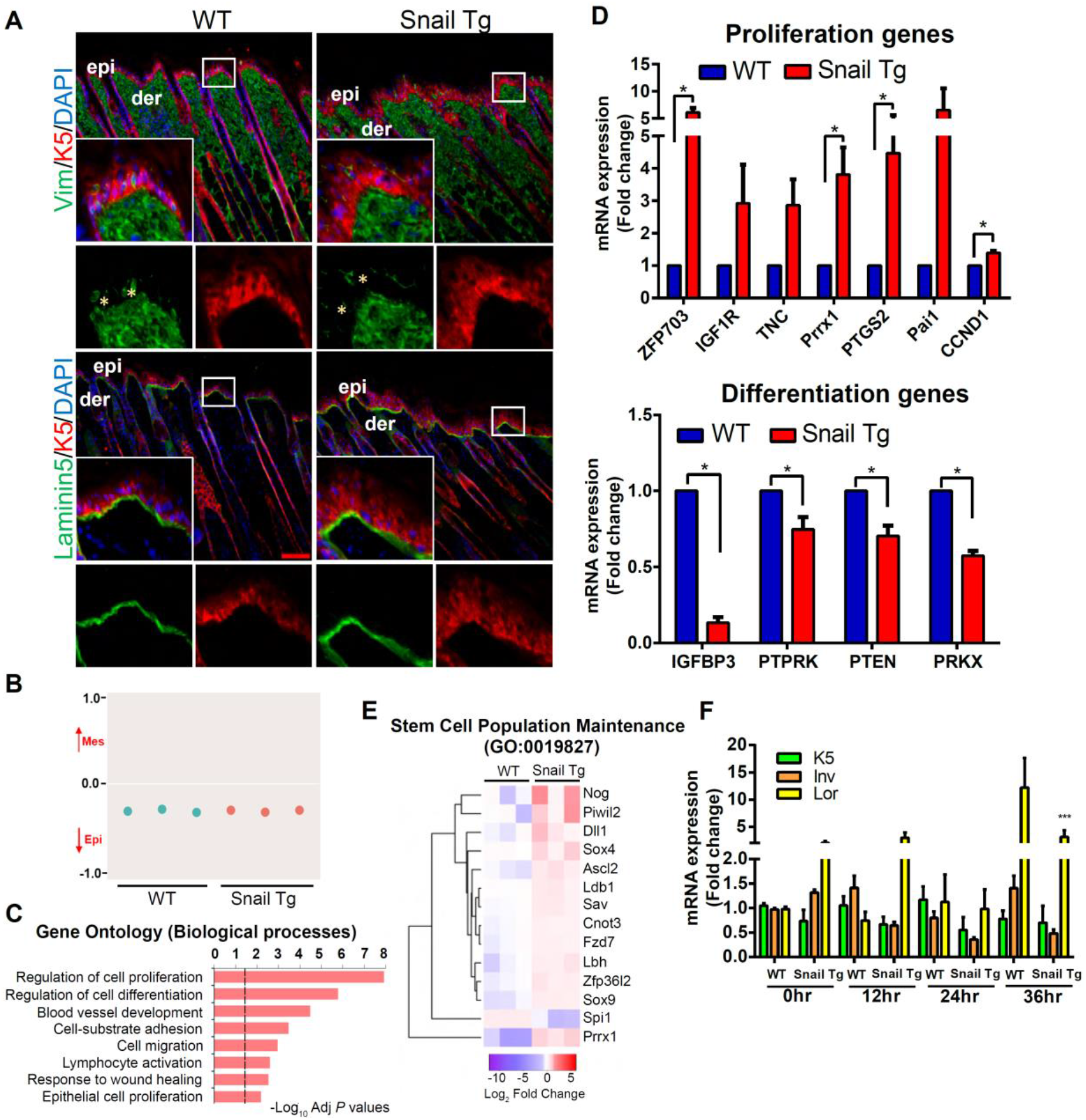
Overexpression of Snail reinforces the stem/progenitor characteristics in vitro. (A) lmmunofluorescenceassay representing intact compartmentalization of cells in the skin. Top row: Vimentin (green) to mark the mesenchymal cells, Keratin 5 (red) to mark the basal epidermal keratinocytes and outer root sheath of hair follicles in wild type (WT) and Snail transgenic (Tg) skin. Bottom row: Laminin 5 (green) marks the basal lamina and keratin 5 (red) marks the basal epidermal keratinocytes. Insets in each panel are magnified views of the boxed areas of the epidermis. Scale bar: 50μm. (B) EMT score calculated using the transcriptome data of WT and Snail Tg keratinocytes. (C) Gene ontology of the differentially expressed genes in Snail Tg keratinocytes. (D) qPCR of proliferation and differentiation associated genes (n=3). (E) Heat map of genes corresponding to the GO term: stem cell population maintenance. (F) qPCR analysis of differentiation potential of WT and Snail transgenic keratinocytes in the calcium switch assay (n=3). Error bars depict± SEM. * p < 0.05, ***p < 0.001. See also Figure S1.

We previously observed that the Snail Tg skin recapitulated many hallmarks of cutaneous squamous cell carcinoma (Du et al., 2010). In particular, there was robust epidermal hyperplasia with an expansion of the basal layer (Du et al., 2010), which harbours the stem/progenitor cells of the epidermis. To determine how Snail is stimulating the expansion of the basal layer of keratinocytes (Du et al., 2010) in an non-EMT fashion, we analysed the transcriptome profile of the transgenic keratinocytes for potential clues. Gene ontology of differentially expressed genes (Figure S1C) revealed that the regulation of cell proliferation and cell differentiation are the top two biological processes that are affected in the Snail Tg keratinocytes (Figure 1C). qPCR analysis confirmed that there was elevated levels of proliferation associated genes and decreased expression of differentiation genes in Snail Tg keratinocytes (Figure 1D). Thus, the underlying expansion of the basal layer of the Snail Tg skin may be accomplished by the hyperproliferation of keratinocytes, and/or the inhibition of differentiation. Interestingly, we found that primary Snail Tg keratinocytes do not display a higher proliferation rate in vitro compared to wild type cells (Figure S1D). This is consistent with our previous report that implicated inflammation as the inducer of epidermal keratinocyte proliferation in vivo (Du et al., 2010). Interestingly, the transcriptome of the transgenic cells indicates that genes involved in stem cell population maintenance are upregulated (Figure 1E and Figure S1E). One prediction that can be made if Snail maintains the stem/progenitor characteristic of basal epidermal keratinocytes, is that there should be resistance to the chemical induction of differentiation. To test this, we inspected the differentiation potential of Snail Tg keratinocytes *in vitro* by the classical calcium switch assay (Bikle et al., 2012). Differentiation genes such as *involucrin* and *loricrin* are lower in the Snail Tg keratinocytes compared to WT cells (Figure 1F and Figure S1F). This is consistent with our previous study which reports that the Snail Tg epidermis exhibits a spatial delay in the differentiation process (Jamora et al., 2005). In total, these results reveal a novel non EMT function of Snail in maintaining stem/progenitor characteristics in epidermal keratinocytes of the basal layer.

### Epidermal Snail expression expands the epidermal stem/progenitor cell population *in vivo*

To investigate whether the epidermal hyperplasia in the Snail Tg skin (Du et al., 2010) is due to the expanded number of epidermal stem/progenitors cells, we analysed two widely utilized markers of these cells – CD49f (Krebsbach and Villa-Diaz, 2017; Schober and Fuchs, 2011; Ye et al., 2017) and p63 (Blanpain and Fuchs, 2007; Koster et al., 2005). Immunohistochemical analysis of these markers showed an increase in the Snail Tg epidermis compared to its WT counterpart (Figure 2A). These results were intriguing since the basal layer (marked by Keratin 5) is the exclusive home of the epidermal stem/progenitor cells and their delamination from this layer to the suprabasal layer is generally considered a signal to lose their stemness and enter the terminal differentiation program (Fuchs, 2008). We found that there is a substantial increase in the number of p63 positive cells in the suprabasal layers of the Snail Tg epidermis (Figure S2A). In addition, generic markers of stem cells such as *Oct4, Sox2* and *Nanog* were likewise upregulated in the Snail Tg skin (Figure S2B). Furthermore, ultrastructural analysis of the suprabasal keratinocytes of the epidermis in the Snail Tg skin revealed pleomorphic nuclei with diffused heterochromatin and multiple nucleoli, which are morphological features of stem cells (Figure 2B).

**Figure 2.**
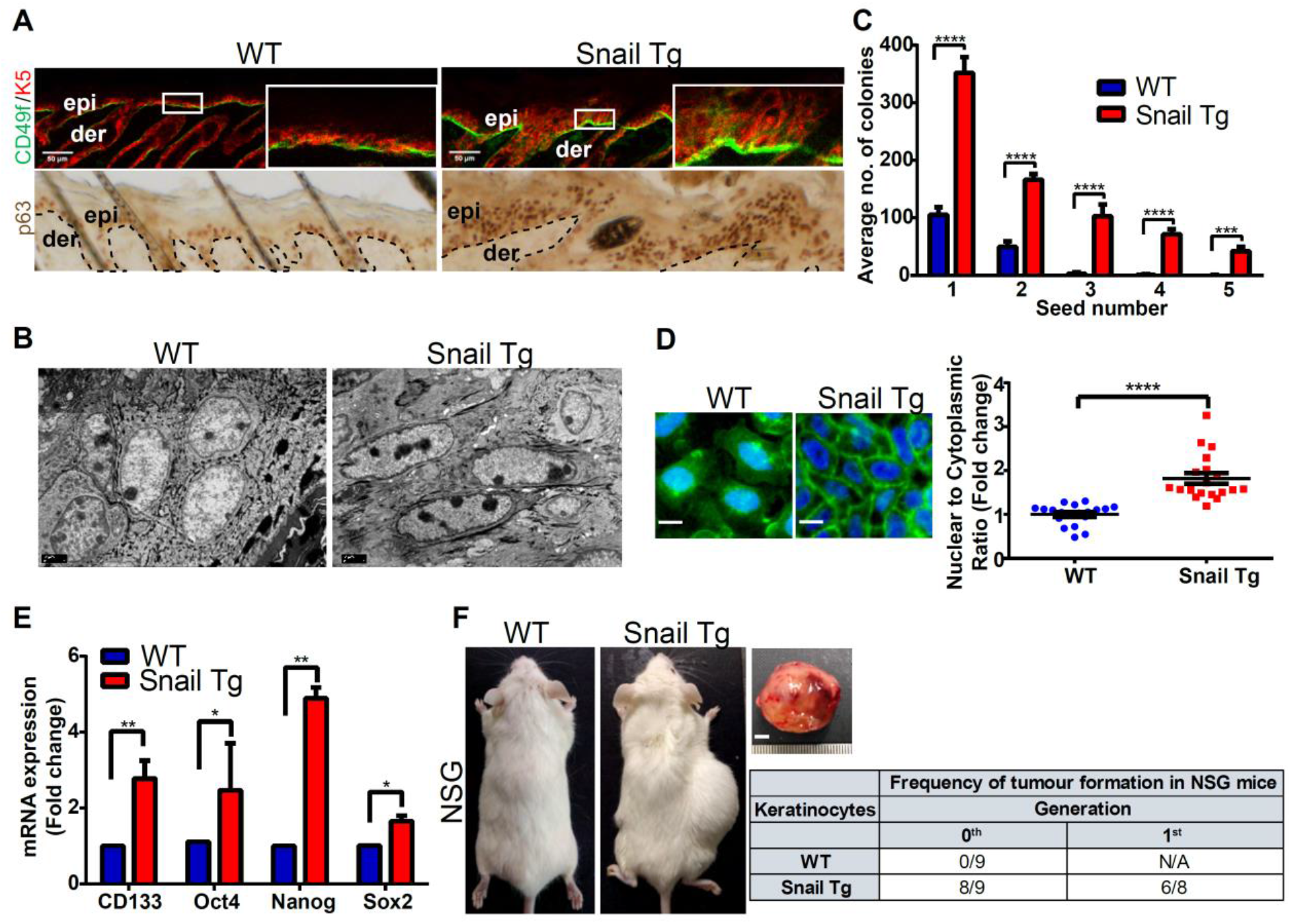
Epidermal Snail expression expands the epidermal stem/progenitor cell population in vivo. (A) lmmunohistochem istry of CD49f and p63 marking the stem/progenitor cells in WT and Snail Tg skin. Dotted line represents the basement membrane, which separates the epidermis (epi) from the underlying dermis (der). Scale bar: 50μm (B) Ultrastructure analysis demonstrating increased number of nucleoli and dispersed chromatin in the nucleoplasm of Snail Tg suprabasal keratinocytes compared to its WT counterpart (n=2). Scale bar: 2μm (C) Colony formation assay performed on WT and Snail Tg keratinocytes. 1000 cells were plated per well for each seeding and number of colonies were quantified seven days post seeding (n=3/seeding) (D) Analysis of nuclear to cytoplasmic ratio in WT and Snail Tg epidermal keratinocytes. Left panels: lmmunofluorescence of WGA to mark the cell membrane and DAPI to mark the nucleus. Scale bar: 5μm Right panel: Quantification showing higher nuclear to cytoplasmic ratio of Snail Tg keratinocytes compared to WT cells. (E) WT (blue) and Snail Tg (red) keratinocytes analysed for the stem cell genes CD133, Oct4, Sox2, and Nanog by qPCR (n=3). (F) Tumor forming capacity of WT and Snail Tg keratinocytes injected in NSG animals. Image shows tumour formed by the Snail Tg keratinocytes at the flank region two months post injection. Scale bar: 5mm. The table summarizes the number of tumours formed by WT and Snail Tg cells in the initial injection and the 1st generation derived from cells from the tumor developed in the initial injection. Error bars depict± SEM. * P < 0.05, ** P < 0.01, *** P < 0.001 and **** p < 0.0001. See also Figure S2.

Given the expansion of the epidermal stem/progenitor cells found *in vivo*, we investigated whether this phenotype is a cell autonomous effect of *Snail* overexpression or secondary to inflammation (Du et al., 2010). To probe the cell autonomous function of *Snail*, we cultured primary keratinocytes from WT and Snail Tg skin and analysed various characteristics of stemness. In colony forming assays, which scores for the self-renewal capacity of stem cells, Snail Tg keratinocytes exhibited a higher number of colonies over several seedings compared to WT cells (Figure 2C). Other features of CSCs such as increased nuclear to cytoplasmic ratio (Yang et al., 2020) (Figure 2D), and expression of stem cell genes (Figure 2E) are higher in Snail Tg keratinocyte relative to its WT counterpart. These observations are consistent with reports of a strong positive correlation between Snail expression and the undifferentiated state of cancer cells (Fan et al., 2012a; Ma et al., 2017). Our *in vitro* findings were further validated using a xenograft assay, which is the gold standard in defining a cell as a stem cell. We grafted 0.25×10^6^ Snail Tg keratinocytes in NSG animals with WT cells as control. Two months post grafting, ~90% of mice formed tumors with Snail Tg keratinocytes compared to 0% with control cells (Figure 2F). Interestingly, when we serially grafted the cells isolated from the first graft (1^st^ generation) of Snail Tg tumors, we observed that ~75% of the grafts formed 2nd generation tumors. These results suggest that *Snail* overexpression not only upregulates markers associated with stem/progenitor cells, but it also endows them stem cell capabilities.

### Snail mediated stem/progenitor cell maintenance is dependent upon STAT3 activation

While many studies have firmly established an association of Snail expression with heightened cancer stem like properties (De Craene et al., 2014; Hojo et al., 2018a), the mechanism underlying this correlation nevertheless remains elusive. An essential component that maintains the stemness of a variety of cells is the active form of STAT3 (phosphorylated STAT3)(Galoczova et al., 2018; Raz et al., 1999). Previously, we had shown that there is increased activation of STAT3 in the Snail Tg compared to the WT epidermis (Du et al., 2010). The activation of STAT3 was also observed in cultured Snail Tg keratinocytes (Figure 3A-B and Figure S3A-B). To determine the functional relevance of this activated STAT3, we treated WT and Snail Tg keratinocytes with BP-1-102, a small molecule inhibitor, which inhibits the phosphorylation, dimerization, and subsequent nuclear translocation of STAT3 (Zhang et al., 2012). We found that inhibition of STAT3 activity by ~50% in Snail transgenic keratinocytes (Figure S3C-D) substantially suppressed their self-renewal capacity (Figure 3C) and stem cell associated genes (Figure 3D). In addition, the increased nuclear to cytoplasmic ratio (NCR) and the expression of the cancer stem cell marker CD44 (Figure 3E-F and Figure S3E) were significantly reduced to near WT levels upon STAT3 inhibition.

**Figure 3.**
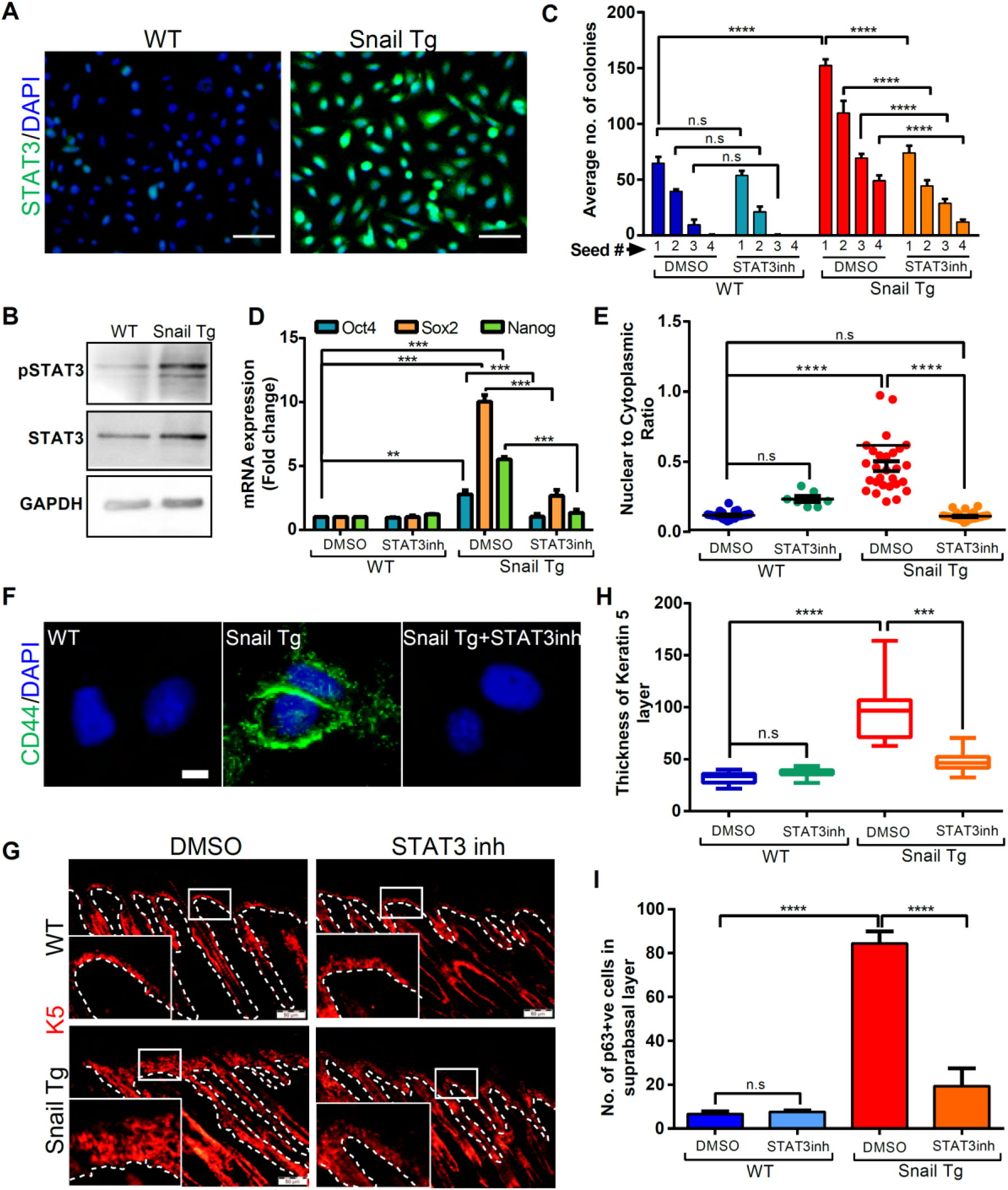
Snail mediated stem/progenitor cell maintenance is dependent upon STAT3 activation. (A) lmmunofluorescence of total STAT3 (green) and DAPI (blue) on WT and Snail Tg keratinocytes. Scale bar= 10μm. (B) Western blot for phosphorylated STAT3 (pSTAT3) and total STAT3 in WT and Snail Tg keratinocytes. GAPDH was used as loading control. (C) Average number of colonies formed by WT and Snail Tg keratinocytes in the presence or absence of STAT3 inhibitor or vehicle control, DMSO. n=3 for each seeding. (D) qPCR analysis of stem cell gene expression in WT and Snail Tg keratinocytes upon STAT3 inhibition (n=3). (E) Nuclear to cytoplasmic ratio of WT and Snail Tg keratinocytes upon STAT3 inhibition. (F) lmmunofluorescenceof stem cell marker CD44 in green in WT, Snail Tg, and Snail Tg+STAT3inh cells. Scale bar: 2μm. (G) lmmunofluorescence panels showing stem/progenitor layer of the epidermis marked by keratin 5 in red on WT and Snail Tg skin upon STAT3 inhibition. Dotted line represents the basement membrane, which separates the epidermis (epi) from the underlying dermis (der). Insets in each panel are magnified views of the boxed areas of the epidermis. Scale bar: S0μm (H) Quantification of thickness of the stem/progenitor layer in the WT and Snail Tg skin post STAT3 inhibition, n=2 animals each. (I) Quantification of the number of p63 positive cells in the suprabasal layer of the WT and Snail Tg epidermis after STAT3 inhibition. Error bars depict± SEM. **p < 0.01, ***p < 0.001, P<.000a1nd n.s=not significant. See also Figure S3.

We extended this study to understand if STAT3 activation was essential for Snail’s impact on keratinocyte stemness *in vivo*. The STAT3 inhibitor was applied topically on the back skin of newborn WT and Tg animals every two days up to day 7. Analysis of the skin revealed that the increase in the number of layers positive for keratin 5 in the Snail Tg epidermis was significantly reduced upon STAT3 inhibition (Figure 3G-H). Further we found that the reduction of the thickened basal layer correlated with a decrease in the number of p63+ stem/progenitor cells (Figure 3I and S3F). Collectively, both *in vitro* and *in vivo* studies indicate that the activation of STAT3 is crucial for Snail mediated stem/progenitor characteristics of epidermal keratinocytes.

### Mindin secreted by Snail Tg keratinocytes activates STAT3 in an autocrine fashion

We further sought to elucidate the molecular pathway linking Snail with STAT3 activation. STAT3 activation is regulated via receptor tyrosine kinases (Grandis et al., 1998) or by non-receptor tyrosine kinases (Bowman et al., 2000), which are stimulated by multiple extracellular ligands (Rébé et al., 2013). We therefore postulated that keratinocytes expressing Snail secrete proteins to the extracellular milieu capable of activating STAT3. To test this hypothesis, we treated primary keratinocytes with conditioned media obtained from cultured WT or Snail Tg keratinocytes and assessed the levels of STAT3 phosphorylation. Snail Tg conditioned media increased the levels of activated STAT3 compared to wild type conditioned media (Figure 4A and S4A). We next embarked upon identifying the specific component/s in the Snail Tg conditioned media that is responsible for this effect. Interestingly, we observed increased levels of secreted Mindin in non-permeabilized Snail Tg skin compared to WT tissue (Figure S4B). Likewise, Mindin RNA is upregulated in the Snail Tg keratinocytes compared to the WT cells (Figure 4B). Mindin has attracted recent attention due to its upregulation in numerous carcinomas (Jin et al., 2017; Lucarelli et al., 2013a; Ni et al., 2019; Simon et al., 2007; Zhang et al., 2015) which underlies its use as a diagnostic biomarker. Consistent with a role in mediating keratinocyte stemness, the epidermal hyperplasia found in the Snail Tg skin is reduced to almost WT levels when Mindin was genetically ablated (Figure 4C and Figure S4C). Moreover, we observed that the expansion of the stem/progenitor population in the epidermis of Snail Tg skin, is reduced in the absence of Mindin (Figure 4C). Further, we examined if Mindin is the link between Snail and STAT3 activation. In Snail Tg/Mindin KO skin, there was a significant reduction in the levels of activated STAT3 compared to the Snail Tg skin (Figure 4C). Concomitant with the in vivo observations, Snail Tg keratinocytes lacking Mindin showed reduced self-renewal capacity (Figure 4D), decreased expression of stem cell genes (Figure 4E), and a lower nuclear to cytoplasmic ratio (Figure 4F). Moreover, the level of activated STAT3 in the Snail Tg/Mindin KO keratinocytes was lower compared to Snail Tg cells (Figure S4D).

**Figure 4.**
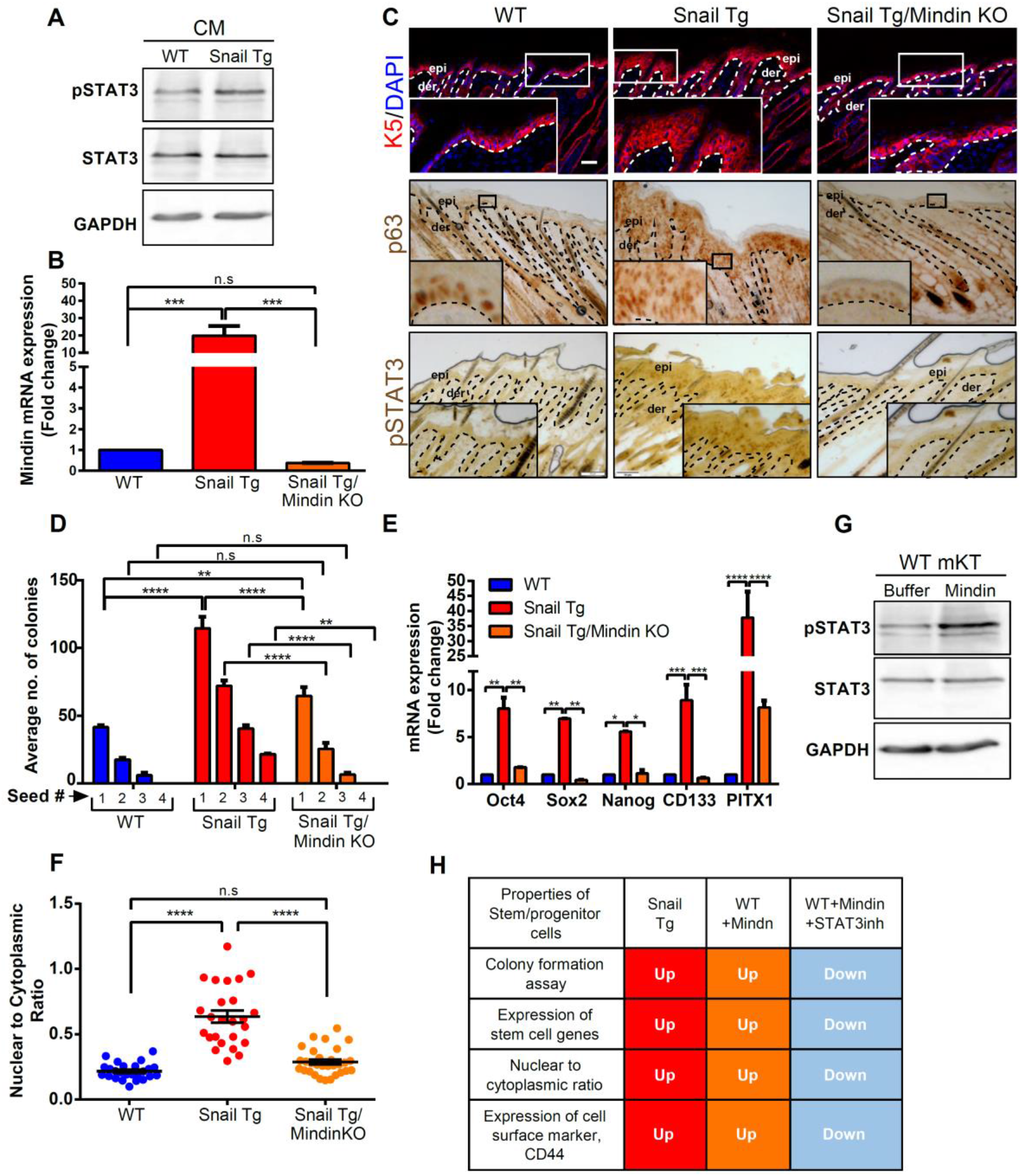
Secreted Mindin by Snail Tg keratinocytes activates STAT3 in an autocrine fashion. (A) Western blot for pSTAT3 and total STAT3 in WT cells treated with WT and Snail Tg epidermal keratinocyte conditioned media (CM). GAPDH was used as loading control (n=3). (B) qPCR analysis showing the Mindin mRNA levels in WT, Snail Tg, and Snail Tg/Mindin KO epidermal keratinocytes (n=3). (C) Top row: lmmunofluorescence of keratin 5 (KS) in red to mark the stem/progenitorlayer of the epidermis in WT, Snail Tg, and Snail Tg/Mindin KO skin. Scale bar: 30μm. Middle row: lmmunohistochemistry of keratinocyte stem cell marker, p63 in brown on WT, Snail Tg, and Snail Tg/Mindin KO skin. Bottom row: lmmunohistochemistry of pSTAT3 in brown on WT, Snail Tg and Snail Tg/Mindin KO skin. Dotted line represents the basement membrane, which separates the epidermis (epi) from the underlying dermis (der). Scale bar: 50μm. (D) Average number of colonies formed by WT, Snail Tg, and Snail Tg/Mindin KO keratinocytes (n=3). (E) mRNA expression of stem cell genes Oct4, Sox2, Nanog, CD133, and PITX1 in wild type Snail Tg and Snail Tg/Mindin KO keratinocytes (n=3). (F) Quantification of the nuclear to cytoplasmic ratio of WT, Snail Tg, and Snail Tg/Mindin KO keratinocytes. (G) Western blot for pSTAT3 and total STAT3 in WT keratinocytes treated with recombinant Mindin and buffer as control. GAPDH was used as loading control (n=3). (H) Table summarizing the stem/progenitorproperties displayed by WT, Snail Tg, WT+ Mindin, and WT+Mindin+STAT3inh keratinocytes. ‘Up’ depicts an increase compared to WT and ‘Down’ depicts a decrease compared to Snail Tg keratinocytes. Error bars depict ± SEM. * **P**< 0.05, ** P < 0.01, *** P < 0.001, **** p < 0.0001 and n.s=not significant. See also Figure S4.

To test the sufficiency of Mindin to activate STAT3, we treated WT keratinocytes with recombinant Mindin and analysed the levels of phosphorylated protein. We observed that Mindin treatment resulted in elevated levels of activated STAT3 (Figure 4G, Figure S4E-F). We further investigated if Mindin is itself sufficient to enhance the stem/progenitor characteristics. Mindin treated keratinocytes exhibited an increased colony forming capacity (Figure S4G), a higher NCR (Figure S3H), and elevated expression of stem cell markers (Figure S4I-J) in a STAT3-dependent manner. In summary, Mindin is sufficient to endow epidermal keratinocytes with stem/progenitor characteristics through the activation of STAT3 (Figure 4H).

### Src Family of Kinases mediates Mindin dependent activation of STAT3

We then investigated the kinase responsible for phosphorylating STAT3 downstream of Mindin signalling. Mindin is reported to be ligand for integrins on immune cells which in turn activates STAT3 via Src Family of Kinases (SFKs) (Jia et al., 2005a; Xue et al., 2010). To test whether this same pathway is active in epidermal keratinocytes, we assessed the levels of activated (phosphorylated) Src in Snail Tg vs. WT keratinocytes with a phospho-pan Src antibody (Figure 5A and Figure S5A). Snail Tg keratinocytes exhibited a two-fold increase in activated Src compared to its WT counterpart. To examine whether Mindin was sufficient to activate Src, we treated WT keratinocytes with recombinant protein. Mindin treated cells exhibited a three-fold increase in Src activation (Figure 5B and Figure S5B).

**Figure 5.**
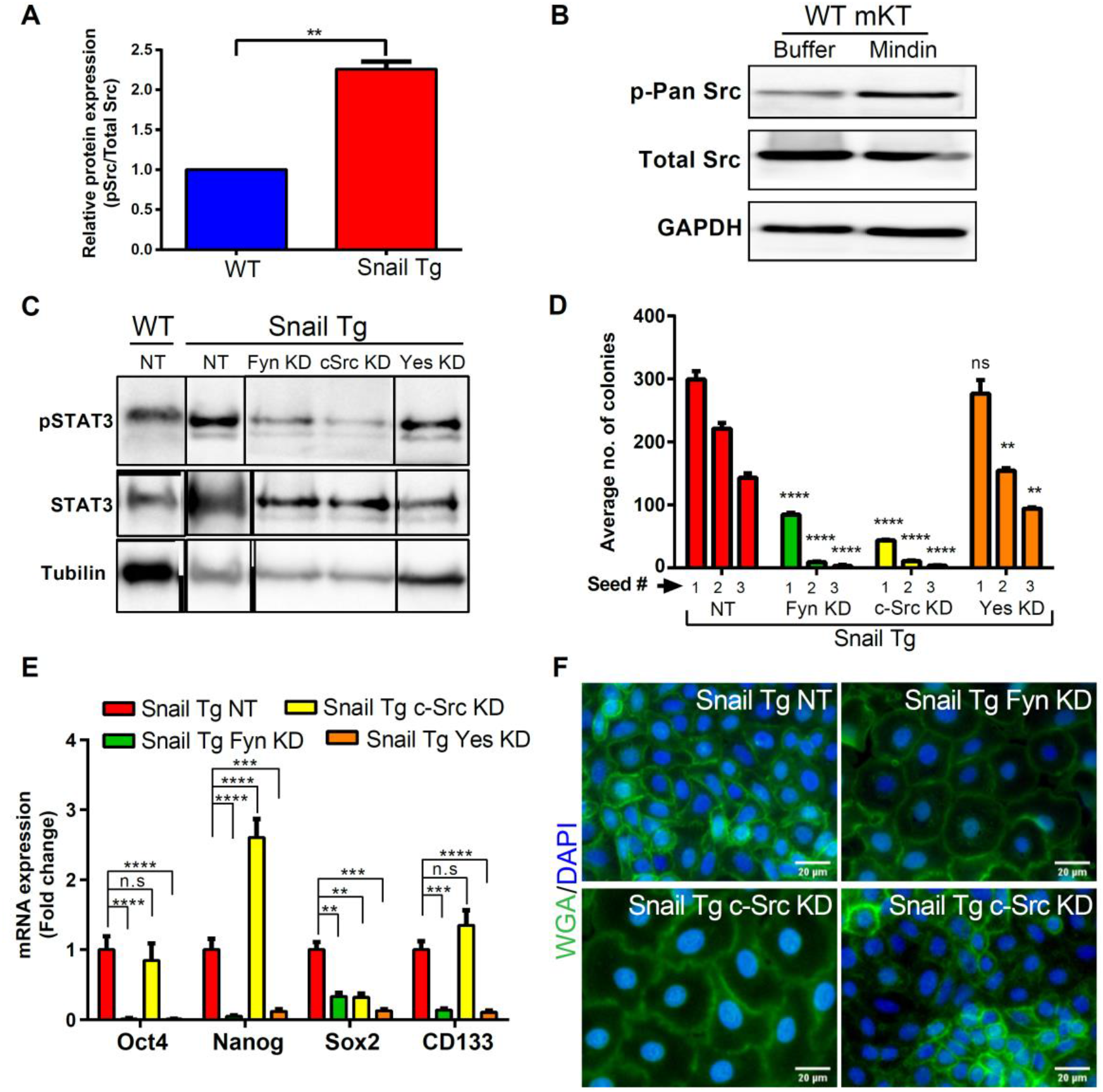
Src Family of Kinases mediate the Mindin dependent activation of STAT3. (A) Quantification of western blot analysis of phospho pan Src to total Src in WT and Snail Tg keratinocytes (n=3). (B) Western blot for phospho pan Src and total Src in WT keratinoc ytes treated with Mindin protein and its buffer control. GAPOH was used as the loading control (n=3). (C) Western blot for pSTAT3 and total STAT3 in WT and Snail Tg keratinocytes with non-targeting (NT) control and Snail Tg keratinocytes knocked down (KO) individually for Fyn, c-Src and Yes. Gamma tubulin was used as the loading control. (D) Average number of colonies formed by Snail Tg keratinocytes knocked down for Fyn, c-Src and Yes with NT as control (n=3). (E) mRNA express ion of stem cell genes in the Snail Tg keratinocytesindividually knocked down for Fyn, c-Src and Yes with NT as control (n=3). (F) lmmunofluorescence of WGA in green marks the plasma membrane and OAPI in blue marks the nucleus. Nuclear to cytoplasmic ratio has reduced in Snail Tg cells with Fyn and c-Src KO compared to the Snail Tg NT and Yes KO cells. Scale bar: 20μm. Error bars depict± SEM. * P < 0.05, ** P < 0.01, *** P < 0.001, ****p < 0.0001 and n.s=not significant. See also Figure S5.

A broad-spectrum pharmacological inhibitor of SFKs led to a reduction in phosphorylated STAT3 levels in Snail Tg keratinocytes (Figure S5C). We then explored which specific SFK member(s) is responsible for Mindin mediated STAT3 activation using a knockdown approach. shRNA mediated knockdown was performed for each of the three ubiquitously expressed SFK members (*c-Src, Fyn and Yes*) in Snail Tg keratinocytes (Figure S5D). Knocking down either *Fyn or c-Src* resulted in a significant reduction in the levels of phosphorylated STAT3 (Figure 5C and Figure S5E). Furthermore, to functionally examine the effect of these knockdowns, we assessed their impact on the colony forming capacity of Snail Tg cells. Reduced levels of either *Fyn or c-Src* resulted in fewer colonies, while the *Yes* knockdown showed a mild effect (Figure 5D). Interestingly cells with lower levels of *Fyn or Yes* exhibited a reduced expression of stem cell genes (Figure 5E). Finally, we characterised SFK on the morphological parameters of keratinocyte stemness and found that *Fyn* and *c-Src* knock down caused a decrease in the nuclear to cytoplasmic ratio while the *Yes* knock down had no effect (Figure 5F and Figure S5F). Overall, these results suggest that Mindin-STAT3 mediated stem/progenitor properties is via activation of Src kinases. Notably, Fyn appears to have the broadest effect on the stem cell characteristics we analysed (Figure S5G).

### Mindin is required for the maintenance of stem/progenitor characteristics in human cutaneous squamous cell carcinoma

Although Mindin has been proposed to be a biomarker for certain cancers such as ovarian (Simon et al., 2007), prostate (Lucarelli et al., 2013b), and lung adenocarcinoma (Yuan et al., 2017), the precise role of Mindin in skin cancers remains elusive. Therefore, we sought to determine if Mindin mediated stemness observed in Snail Tg mouse keratinocytes is operational in human skin cancers. We first inspected the expression pattern of Mindin in various cancers and found it to be upregulated in 12 out of the 18 different cancers that were analysed (Figure S6A). In addition, a cell line (A388) derived from a patient with cutaneous squamous cell carcinoma (cSCC) (Conway et al., 1992) that expresses higher levels of Snail (Figure S6B), also exhibited higher levels of Mindin mRNA (Figure 6A). We proceeded to investigate whether the increase in Mindin levels observed in the cSCC cell has a functional role in maintaining stem/progenitor characteristics. We utilized the CRISPR/Cas9 system to knockdown Mindin in cSCC cells (cSCC-KD). cSCC cells that were selected for Mindin KD with puromycin contained a 50% reduction in the mRNA levels of Mindin compared to control cells (Figure S6C). cSCC-KD cells displayed a reduction in the expression levels of stem cell genes (Figure 6B), self-renewal capacity (Figure 6C) and nuclear to cytoplasmic ratio (Figure 6D). Moreover, the trend observed in all these properties could be rescued upon treatment of cSCC-KD cells with recombinant Mindin, suggesting that the phenomenon is not an off-target effect of gene editing. Furthermore, we investigated whether the Mindin mediated Src-STAT3 signalling module is active in human cSCC cells. cSCC-KD cells exhibited reduced p-STAT3 (Figure 6E) and p-Src (Figure 6F) levels compared to its control (Figure 6E). Moreover, the reduction in SFK and STAT3 activation in cSCC-KD cells was reversed upon the treatment with recombinant Mindin. Supporting these results, the stem cell markers associated with the cSCC cells also decreased in the KD cells (Figure 6G-H).

**Figure 6.**
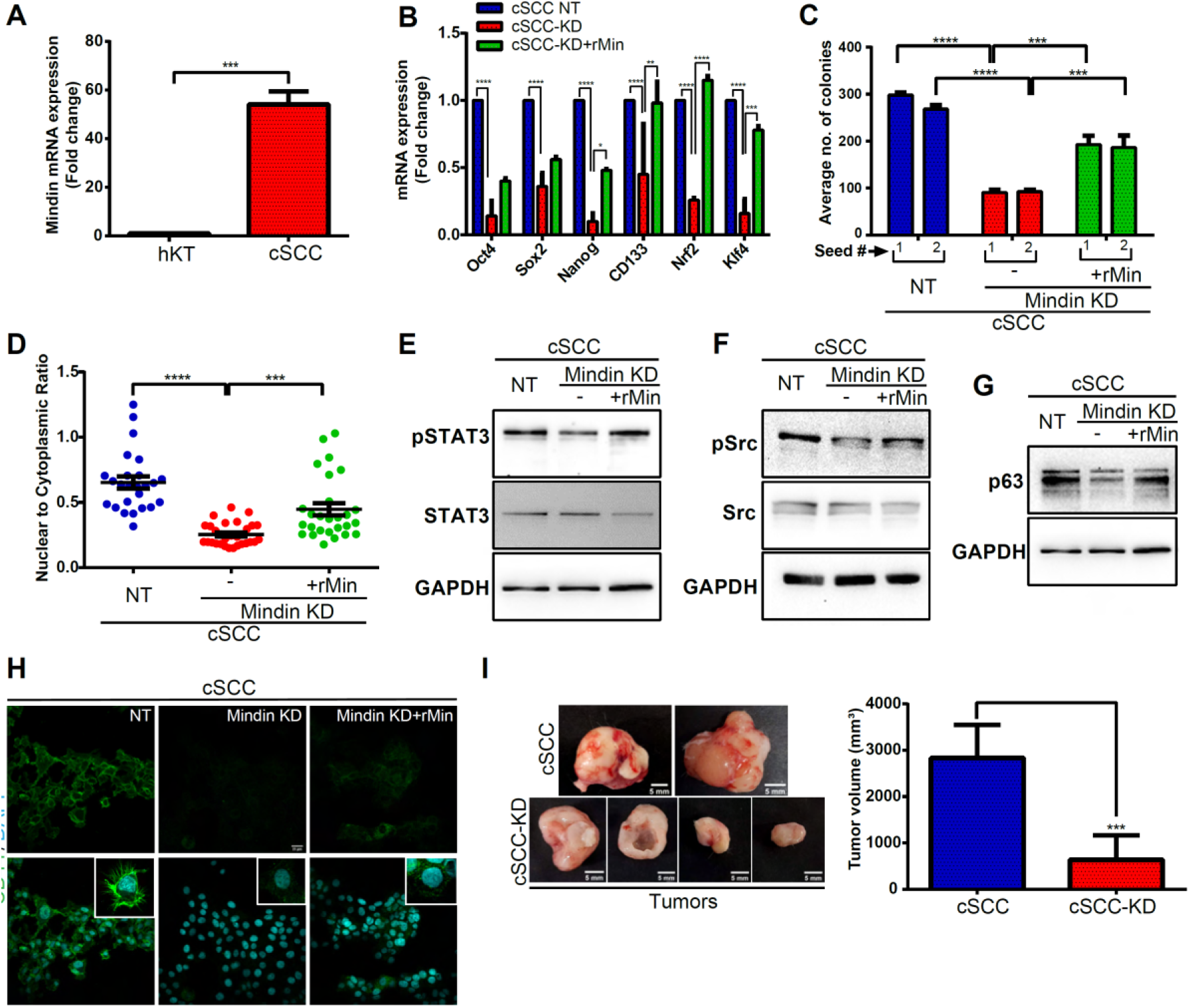
Mindin is required for the maintenance of stem/progenitor characteristics in cutaneous squamous cell carcinoma cells. (A) qPCR for Mindin mRNA in patient derived cutaneous squamous cell carcinoma (cSCC) cell line compared to normal human keratinocytes (n=3). (B) mRNA expression of stem cell genes in cSCC no target (NT) control, cSCC Mindin KO (cSCC-KD) and cSCC-KD treated with recombinant Mindin (n=3). (C) Average number of colonies formed by cSCC NT control cells, cSCC-KD and cSCC-KD cells treated with recombinant Mindin (n=3). (D) Quantification of nuclear to cytoplasmic ratio of cSCC NT control cells, cSCC-KD and cSCC-KD cells treated with recombinant Mindin. (E-G) Western blot analysis for pSTAT3 and total STAT3 (E), pSrc and total Src (F) and p63(G) in cSCC NT control cells, cSCC-KD and cSCC-KD cells treated with recombinant Mindin. (H) lmmunofluorescence of cell surface marker CD44 in green on cSCC NT control cells, cSCC-KD and cSCC-KD cells treated with recombinant Mindin. Scale bar: 25μm (I) Tumors formed by the cSCC cells (top) and by cSCC-KD cells (bottom) in the xenograft assay. Scale bar: 5mm. In the right panel is the tumor volume in mm^3^ of cSCC and cSCC-KD tumors. Error bars depict± SEM. * P < 0.05, ** P < 0.01, *** P < 0.001, *** *p < 0.0001, See also Figure S6.

In xenograft assays, cSCCs formed tumors in 5/5 mice while the cSCC-KD cells formed tumors in 7/10 mice. However, the size of the tumors that formed with the cSCC-KD cells were substantially smaller compared to its parental cSCC cells (Figure 6I). In addition to their size difference, the tumors formed by the cSCC-KD cells unexpectedly appeared to have less vascularization compared to that of the tumors from cSCC cells (Figure 6I). Overall, these results signify that Mindin is a critical player required for the maintenance of stem/progenitor characteristics in human squamous cell carcinoma of skin. In addition, the signalling pathway that Mindin mediates to maintain stem/progenitor characteristics in mouse is conserved in human skin cancer cells.

## Discussion

The predominant model of Snail’s contribution towards tumorigenesis in various carcinomas, invokes its canonical function in promoting an EMT (Fan et al., 2012b; Goossens et al., 2017; Smith et al., 2014; Yang et al., 2017; Zhu et al., 2012). Several studies have established Snail-mediated EMT in either transformed cell lines or cancerous cells that contain multiple genetic aberrations. Surprisingly, we observed that ectopically expressed Snail in primary epidermal keratinocytes, results in the downregulation of E cadherin RNA but not protein(Jamora et al., 2005). The notion of an EMT itself has undergone a renaissance due to the new appreciation of the spectrum of EMT characteristics (Grosse-Wilde et al., 2015; Pastushenko and Blanpain, 2019). Thus the EMT program is no longer considered a binary process with a completely epithelial (E) or mesenchymal (M) phenotype (Grosse-Wilde et al., 2015; Kröger et al., 2019). Instead, an important intermediate called the EM hybrid state, correlates with maximum stem cell characteristics (Bierie et al., 2017). However, computation of the EMT score using the transcriptome profile we generated with transgenic cells revealed a negative score (<0) in both wild type and Snail Tg primary mouse keratinocytes signifying their epithelial nature (Tan et al., 2014). This also suggests that Snail is not sufficient to induce EMT in primary cells and requires an additional drive to enter the EMT program that are present in transformed and cancer cell lines.

These results are consistent with studies that have implicated an EMT independent role for Snail in conferring cancer cells with stem cell like traits (De Craene et al., 2014; Goossens et al., 2017; Ma et al., 2017; Mani et al., 2009; Zhou et al., 2014). Although this association is well-recognized, the mechanistic basis underlying this connection has thus far remained elusive. Our findings indicate that *Snail* expressing keratinocytes utilize Mindin in an autocrine fashion to activate SFK-STAT3 signalling to promote stemness (Figure 7). This study illustrates how carcinomas are a product of factors operating at multiple cellular levels ranging from transcription factors such as Snail, to cytoplasmic proteins (i.e., SFK-STAT3), and extracellular component such as Mindin. Previously, we reported that one mechanism for STAT3 activation in epidermal keratinocytes of the *Snail* transgenic skin is the paracrine signaling of IL-17 released by activated resident γδT-cells (Du et al., 2010). Interestingly, we find that the increase in the number of active γδT-cells in the Snail Tg background is substantially reduced in the absence of Mindin (Figure S7A). This is in line with previous reports demonstrating the role of this protein in recruitment and activation of other immune cells (Jia et al., 2005b). Together, these results suggest that Mindin regulates stemness in a multitiered fashion: it establishes an inflammatory microenvironment that reinforces the intracellular signalling pathway that resists pro-differentiation signals. In addition, the defect in keratinocyte differentiation of Tg cells (Figure 1D and F) may be facilitated by epigenetic modifications since Snail has been previously reported to associate with the PRC2 complex to inhibit epithelial differentiation (Plikus et al., 2015).

**Figure 7.**
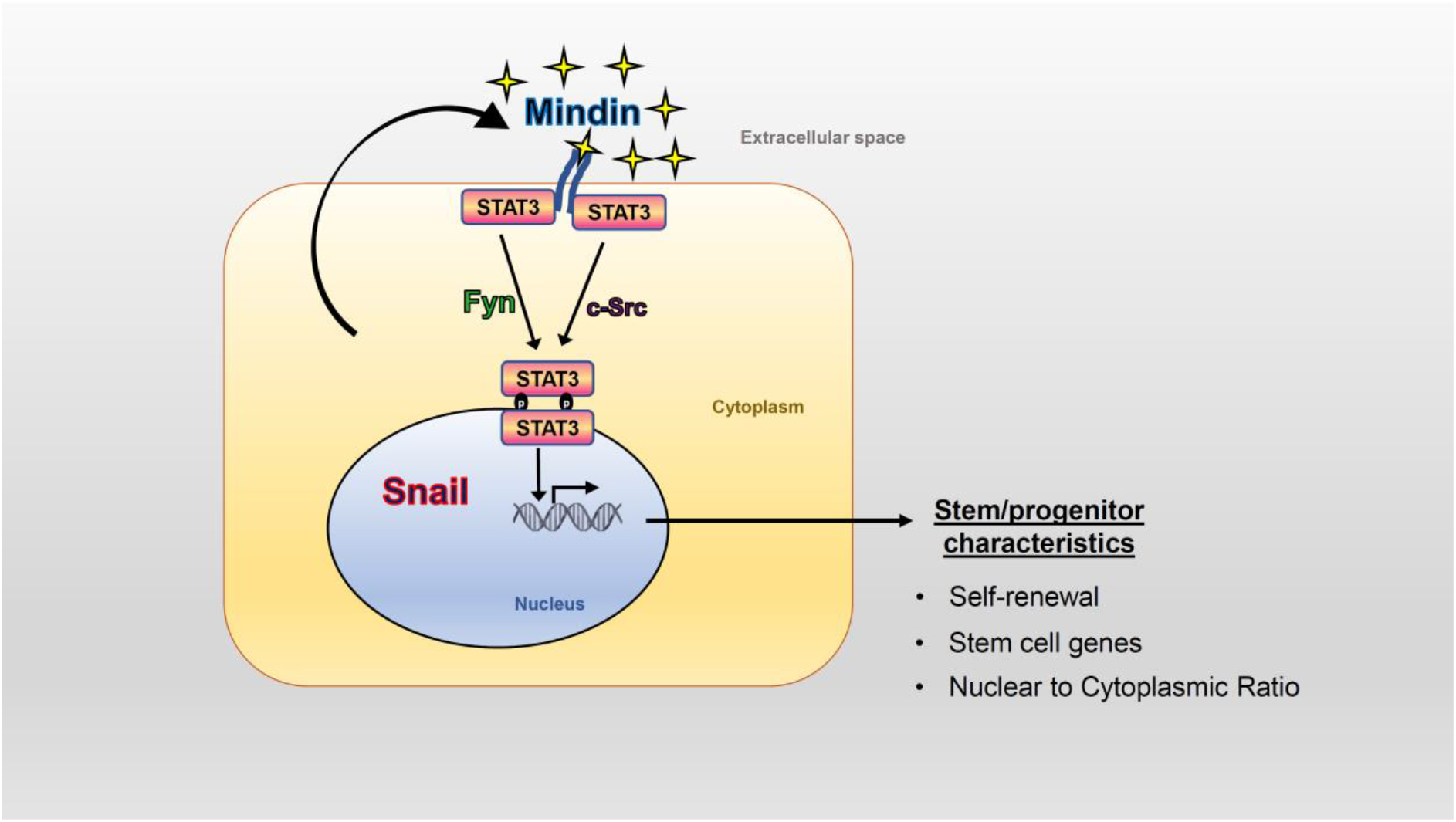
Model indicates that Snail expressing keratinocytes secrete Mindin to the extracellular milieu which in an autocrine fashion activates SFK-STAT3 signalling to promote sternness.

Since we have demonstrated that secreted Mindin is both necessary and sufficient to maintain stemness, one would predict that many cells within its field would be stimulated to manifest stem cell characteristics. Why, then, are the number of cancer stem cells within a tumor so few? There are overlapping possibilities that can account for this apparent contradiction: 1. Only a few cells in the tumor express Snail, so the number of cells secreting Mindin is limited and correlates with the number of cancer stem cells. Interestingly, in aggressive tumors, Snail and Mindin expression is increased along with the number of cancer stem cells (Hojo et al., 2018b; Kang et al., 2020; Zhao et al., 2020); 2. The ECM binding nature of Mindin substantially restricts its diffusion and thus limits the number of cells that is exposed to this matricellular protein; 3. Differential expression of its cognate receptor may limit the number of responding cells. Though the receptor for Mindin on epithelial cells is unknown, it is reported that Mindin is a ligand for integrin αMβ2 and α4β1 on macrophages and neutrophils, respectively (Jia et al., 2005a; Xue et al., 2010). It is noteworthy that these integrins also activate SFK and STAT3 containing signalling pathways (Meng and Lowell, 1998). Moreover, these two integrin pairs are reported to be upregulated in cancers and positively correlate with metastasis and progression (Alday-Parejo et al., 2019; Ganguly et al., 2013; Wu et al., 2021). Identification of the receptor of Mindin on epithelial cells can facilitate expression studies on the heterogenous population of cells within the tumor. This differential expression might explain why only a few cells within a tumor are cancer stem cells.

A notable observation made during the course of this study is that *Snail* expressing keratinocytes exhibit an increase in both activated STAT3 as well as total STAT3 (Figure S3B). The reason for the latter could be a direct binding of Snail on the *STAT3* promoter, which has the canonical E-box sequence for Snail binding (Figure S7B). This would argue that even though Snail is classically considered a transcriptional repressor, it also has the ability to function as a transcriptional activator. This is supported by reports that Snail can transcriptionally upregulate *p15INK4b* (Hu et al., 2010), *MMP15* (Tao et al., 2011) and *Snail2* (Daisuke et al., 2006). This also explains why treatment of keratinocytes with Mindin only results in the upregulation of activated STAT3 without affecting the total amount of this protein (Figure 4G).

Given its role in maintaining stemness, STAT3 has been a focus in cancer therapy and multiple drugs targeting this protein/pathway are currently in clinical trials. However, only a few STAT3 inhibitors are approved by the FDA (Beebe et al., 2018; Zou et al., 2020). One major limitation of targeting STAT3 is that since normal cells require this signalling protein for basic cellular physiology, inhibiting this protein would have many unintended side effects. Therefore, it becomes important to target specific upstream regulators of STAT3 that would allow other physiologically important STAT3 containing signalling pathways to remain operational. In this case, Mindin would be an attractive target as *Mindin* KO mice are viable and fertile and Mindin activates STAT3 in cancer cells. Additionally, in the *Mindin* KO mice, other physiological pathways utilizing STAT3 will still be intact. For instance, the *Snail* Tg/ *Mindin* KO retains STAT3 activation in the basal keratinocytes of the epidermis where it is known to contribute to the proliferative potential of these cells (Figure 4). Finally, the activity of Mindin in both mouse and human squamous cell carcinoma of the skin suggests that this is a conserved process that would be an effective target for therapeutic intervention.

## Materials and Methods

### Animal studies

C57Bl6 and NSG mice were obtained from The Jackson Laboratory (Bar Harbor, Maine) and *Mindin* KO mice was from You-Wen He (Department of Immunology, Duke University Medical University Medical Center) The *K14-Snail* Tg mouse was developed as described earlier (Jamora et al., 2005). The *K14-Snail Tg/Mindin* KO mouse was generated by breeding the *Mindin KO* and *K14-Snail Tg* mice. All mice were maintained and bred at the BLiSC Animal Care and Resource Centre under specific pathogen–free conditions. Mice were sacrificed between P7 and P9 (neonatal) and skin samples were collected for RNA, protein and OCT or paraffin embedding as required. All animal work was approved by the Institutional Animal Ethics Committee in the CJ lab (INS-IAE-2019/06[R1]) and SK lab (NCBS-IAC-2017/05[N]). Experiments on mice followed the norms specified by the Committee for the Purpose of Control and Supervision of Experiments on Animals (Government of India). All experimental work was approved by the Institutional Biosafety Committees of both inStem and NCBS.

### Gene expression

Total RNA was isolated from skin and epidermal biopsies or keratinocytes (either proliferating or differentiated) using TRIzol Reagent or RNAiso Plus (Thermo Fisher Scientific, Takara). 1-2ug of RNA was used to synthesise cDNA using Superscript III or PrimeScript kit (Thermo Fisher Scientific, Takara). Quantitative PCR (qPCR) was done with cDNA equivalent to approximately 100ng of RNA using the Sso Fast 2x master mix (BioRad) in a Bio-Rad CFX384 machine. Actin or GAPDH expression was used as a reference for normalization. The primer sequences used are listed in Supplementary Table 1.

### Transcriptome data analysis

For studying the transcriptome, total RNA was isolated from 6 samples (3 each from WT and Snail Tg keratinocyte samples). The RNA sample QC, NGS library (Illumina) preparation, and sequencing steps were outsourced to a commercial facility. We had received approximately 45-55 million and 80-90 million 150bp paired end reads from WT and Snail Tg samples, respectively. Analysis of the data was done in-house after receiving the raw sequencing reads. For the analysis, established analysis pipeline was used to study our RNA sequencing data. The quality of the raw sequencing reads was checked through FASTQC tool [https://www.bioinformatics.babraham.ac.uk/projects/fastqc/]. The low-quality reads towards both ends were trimmed using “FASTX-TRIMMER” using the parameters as “-f 11-l 140” [http://hannonlab.cshl.edu/fastx_toolkit/index.html]. The trimmed reads (130bp*2) were then aligned to mouse reference genome (mm10, UCSC version) using “HISAT2” (Kim et al., 2019). The “SAM” outputs were converted to “BAM” files, followed by sorting using “Samtools”. Respective “BAM” files were used to generate “Count” matrix using “HTSeq-count” (Anders et al., 2015). The count matrix was used to calculate differentially expressed genes using “DESeq2” R package (Love et al., 2014). We found 1113 genes to be significantly differentially expressed in pair-wise comparison (Adj P values < 0.05). These genes were used for Gene Ontology (GO) enrichment analysis using “DAVID”(Huang et al., 2009).

### Analysis of EMT score of primary keratinocytes

EMT scores were calculated from the transcriptome data for all the samples and computed using the previously published EMT signature and the two-sample Kolmogorov-Smirnov-based method [PMID: 25214461, 28262832]. Gene expression data was mapped to the generic cell line based EMT signature genes (total=218; epithelial=170 and mesenchymal=48) derived using gene expression profiles from tumor and cell lines from various cancers. First, the empirical cumulative distribution function (ECDF) was estimated for Epithelial and Mesenchymal gene sets. Then 2KS test was employed to compute the difference between Mesenchymal ECDF and Epithelial ECDF. The 2KS score was considered as the EMT score. A sample with a positive EMT score and significant P value exhibits a more Mesenchymal phenotype, whereas a negative EMT score and significant P value reflects a more Epithelial phenotype. A sample with insignificant P value is considered intermediate.

### Western blot analysis

The keratinocyte lysates were prepared in RIPA buffer with protease inhibitors (Sigma, #P2714) and sonicated at 4°C. Protein lysate was then mixed with 4X Laemmli sample buffer and heated at 95°C for 3 min before loading on the polyacrylamide gel. This was later transferred onto a nitrocellulose membrane. After the transfer, the membrane was blocked using either 5% BSA (Bovine Serum Albumin) in Tris buffer Saline containing 0.1% Tween (TBST) for phosphorylated proteins or 5% Blotto (Santa Cruz Biotechnology, sc-2325) for 60 min. The blots were probed overnight with the respective primary antibody (Supplementary Table 2). After washing with TBST, the blots were probed with HRP-tagged secondary antibodies and washed again. Signals were detected using Enhanced Chemiluminescence substrate (ECL, Merck) and iBright FL (Thermo) detector. The bands were quantified using Fiji (ImageJ) software and normalized to loading controls.

### Immunostaining and histology

Skin tissues were either fixed in Bouin’s solution, dehydrated, and embedded in paraffin or immediately embedded in OCT (Leica). For staining the OCT sections, 10-15 μm-thick sections were fixed in 4% paraformaldehyde (PFA) before proceeding to the primary antibody staining. Refer the Supplementary Table 2 for the list of primary antibodies used and their respective dilutions. Alexa Fluor 488– or Alexa Fluor 568–labelled secondary antibodies (Jackson ImmunoResearch) were used at a dilution of 1:300. DAPI or Hoechst stain was used to mark nucleus. For paraffin tissue staining, the sections were rehydrated followed by antigen retrieval in 10mM sodium citrate buffer (pH 6). 3% H_2_O_2_ incubation was performed after primary antibody incubation and HRP-labelled secondary antibodies (Jackson ImmunoResearch) was used for IHC. Development was done using DAB substrate (Vector Laboratories; SK4105) to manufacturer’s instructions. For secreted protein staining, the use of detergent was completely avoided to prevent permeabilization, and K5 staining was used as an internal control. Imaging was done on an Olympus IX73 microscope or FV1000 confocal microscope and analysed on the Fiji software.

### Calculation of Nuclear to Cytoplasmic Ratio (NCR)

Primary keratinocytes and A388 cells were cultured on collagen (Millipore) coated coverslips and fixed with 4% PFA. Cells were stained with Wheat Germ Agglutinin-Alexa fluor 488(WGA) (Invitrogen) without permeabilising them. After washes with 1xPBS, cells were stained with DAPI and mounted with 80% glycerol. Images of the cells were obtained on Olympus IX73 microscope. Using the Fiji software, areas of the entire cell (by marking the outline of the cell using WGA as reference) and nucleus (with DAPI as reference) were obtained. Cytoplasmic area was calculated by taking the difference between the cell area and nuclear area. With these measurements, the ratio of nuclear area to cytoplasmic area was calculated.

### Cell culture

Primary epidermal keratinocytes were isolated and cultured as described previously in (Nowak and Fuchs, 2009). The keratinocytes were grown in low calcium (50uM) E-media for most of the assays and for the differentiation assay, E-media containing 1.2mM calcium was used and cultured for 48 hours. Treatment with recombinant Mindin protein and conditioned media was done for 24 hours. Human cutaneous squamous cell carcinoma cell line, A388 provided by Benjamin D. Yu (University of California, San Diego) was cultured in DMEM high glucose media with 10% FBS. All cells were processed as required for RNA and protein extraction or staining.

### Colony formation assay

Self-renewal of cells was measured by colony formation assay as described in (Franken et al., 2006). Briefly, single cells of keratinocytes or cSCC were seeded at a very low-density of 1000 cells/well for each seeding in a 6-well dish and was allowed to form colonies for 7 days. Colonies were counted under a microscope and were trypsinzed for the next seeding.

### Xenograft Assay

Primary keratinocytes (WT and *Snail* Tg) and cSCC cell lines (EGFP ctrl and *Mindin* KD) were cultured and 0.25×10^6^ cells were used per injection for each graft. 1: 1 ratio of cells in media to Matrigel with reduced growth factors (Corning) was prepared on ice. 2 months old male NSG animals (The Jackson Laboratories) which were maintained and bred at the animal facility at the NCBS under specific pathogen–free conditions, were used for this assay. Cells prepared in Matrigel were injected subcutaneously in the right flank region of the mice. Animals were monitored for tumor development. 8 weeks post injection, animals were sacrificed, and the tumors were collected, and measurements were taken. Tumor volume in mm^3^ was calculated using the formula; Volume=[(lengthXwidth^2^)/2]. The tumors were subsequently dissociated into single cells using Dispase (Invitrogen) and trypsin (Sigma) and serially grafted.

### shRNA mediated knock down

All lentivirus production and transduction experiments were done in the BLiSC BSL-2 facility. HEK293T cells well cultured in a 10cm dish till they were 70-80 percent confluent. The cells were then co-transfected with three plasmids- shRNA-pZip-mEf1a (5 μg), psPAX2 (3.75 μg), and pMD2.g (1.25 μg) (total DNA - 10μg) using Lipofectamine LTX reagent (Invitrogen). For transfection, the above-mentioned amounts of DNA were suspended in 500μl plain DMEM to which10μl of plus reagent was added. Meanwhile, 30μl Lipofectamine LTX reagent was dissolved in 500μl plain DMEM in another tube. After incubating 10 min at RT, DNA suspension was added to LTX reagent suspension. The mixture was incubated for 15-20 mins. Fresh DMEM with 10% FBS (5 ml) was added to the HEK293T cells, and the transfection mixture was added dropwise the cells with fresh medium. The media was changed 12-16 hours post transfection. The Lentivirus was cultivated 24hrs, 48hrs and 72hrs post first media change and all collections were pooled. The presence of lentivirus was tested using lenti go-stix. The virus was concentrated in a 100KDa Amicon centrifugation unit (Merck) at 5000g for 15 mins to achieve a 10-fold concentration. The concentrated lentiviral particles were then titrated and added on mouse keratinocytes for infection. 72h post infection %GFP+ cells were estimated. Titre that gave 50-60% transduction efficiency was then used for further downstream experiments. For all downstream experiments 72h post infection cells were exposed to puromycin selection (1μg/ml) for 4 days. shRNA constructs against mouse c-Src, Fyn and Yes was procured from Transomics in vector pZIP-mEf1a. For the sequences refer Supplementary Table 3.

### CRISPR/Cas9 mediated *Mindin* knockdown in cSCC cells

CRISPR/Cas9 tool was used to knockdown *Mindin* in cSCC cells (cSCC-KD). The Lentiviral backbone transfer plasmid p-lentiCRISPR - EGFP sgRNA 1 (Addgene Cat #51760) was used to express human codon-optimized Cas9 protein. The puromycin resistance in this plasmid utilizes EFS promoter and an EGFP targeting synthetic single-guide RNA (sgRNA) element from U6 promoter. This plasmid was used as the negative control in our study. The viral packaging plasmid psPAX2 (Addgene Cat# 12260) and viral envelope plasmid pMD2.G (Addgene Cat# 12259) were purchased from Addgene USA.

#### Plasmid engineering

The EGFP sgRNA 1 sequence of p-lentiCRISPR - EGFP sgRNA 1 plasmid has been replaced by customized *Mindin* specific single-guide RNA (sgRNA) **M226R**<u>*CC***G**</u>CGCATAGCTCCGACTACAGC (226 nucleotide regions of NM_012445.4:44-1039 Homo sapiens spondin 2) using Quick change mutagenesis with h-M226RgRNA RP (CGCATAGCTCCGACTACAGCGGTGTTTCGTCCTTTCCAC) and h-M226RgRNA FP (GCTGTAGTCGGAGCTATGCGGTTTTAGAGCTAGAAATAGCAAGTTAAAATAAG). The plasmid was further confirmed with sequencing. This transfer vector plasmid is named as p-lentiCRISPR-M226R sgRNA.

#### Lentivirus production and Transduction

The Lentivirus production work was performed in BSL2 facility following the assigned guidelines. ~4 million HEK293T cells were seeded to 10 cm dish. After a day, the cells were co-transfected with transfer plasmid (6μg), packaging plasmid psPAX2 (3μg) and envelope plasmid pMD2.G (3μg) using Lipofectamine 2000 Transfection Reagent (Invitrogen). Post 18 hr, transfection media was changed with fresh virus production media (DMEM+10% FBS). After 48 and 72 hours, the 10 ml media supernatant having lentiviral particles was collected and filtered with 0.45μm membrane. The 20 ml media was further concentrated to 40 times (500 ul). The concentrated viral particles were frozen in 25 microliters aliquots in −80-degree deep freezers. Both M226R and negative control lentiviral particles were used further to infect the cSCC cells. The cSCC cells (0.3 million) were seeded into in 6 well plate. The cells were treated with various dilutions of lentiviral particles (0, 50, 100, 200, 400 and 500). After 24hr post infection the media was changed with fresh media containing puromycin (1μg/ml). The appearance of resistance foci was observed in various dilutions as compared to zero dilution treatment. The 400^th^ dilution provided single isolated foci which were used for the experiments. The puromycin selected cells were cultured in DMEM media containing 10%FBS and cells were collected for RNA extraction and validation of the knockdown.

### Electron microscopy

Skin samples from P7 WT and *Snail* Tg pups were fixed in 2% glutaraldehyde, 4% formaldehyde in 0.05 M sodium cacodylate, and 2 mM calcium chloride and then embedded in EPON resin and processed. Slices were imaged using the MERLIN Compact VP Scanning Electron Microscope in the BLiSC EM facility.

### Statistics

Comparisons of 2 groups were done using a 1-tailed, paired Student’s *t* test or a 1-tailed Mann-Whitney *U* test. One-way ANOVA followed by Tukey’s post hoc analysis and Two-way ANOVA was used for multiple group comparisons. GraphPad Prism 6 (GraphPad Software) was used for all statistical analyses. Data represent the mean ± SEM. *P* values of less than 0.05 were considered significant.

## Acknowledgements

The authors would like to thank Satyajit Mayor, Maneesha Inamdar, Subhasri Ghosh and members of the Jamora laboratory for their critical review of the work and insightful discussions. The authors also thank Drs. Deepak Arya, Sangeetha Raajkamal, and Yogesh Chandra, and Gaurav Kansagara for their technical assistance. This work was supported by core funds from the Institute for Stem Cell Science and Regenerative Medicine (inStem), Bellary Road, Bangalore, India and grants from the Department of Biotechnology of the Government of India (BT/PR8738/AGR/36/770/2013) and (BT/PR32539/BRB/10/1814/2019), the National Institute of Arthritis and Musculoskeletal and Skin Diseases (NIAMS), NIH (5R01AR053185-03); and the American Cancer Society (15457-RSG-08-164-01-DDC), and by a Hellman Faculty Fellowship to CJ; Work in the SK lab is supported by National Center for Biological Sciences, Tata Institute for Fundamental Research (NCBS-TIFR) planned funds. RKZ was supported by a Junior Research Fellowship from the Department of Biotechnology (DBT/JRF/13/AL/486). Animal studies were partially supported by the National Mouse Research Resource (NaMoR) grant BT/PR5981/MED/31/181/2012;2013-2016;2018 and 102/IFD/SAN/5003/2017-2018 from the Department of Biotechnology. We thank the staff of the BLiSC Animal Care and Resource Centre for assistance with animal husbandry, the BLiSC Central Imaging and Flow Cytometry Facility for help with image acquisition, and the BLiSC Electron Microscopy facility for ultrastructural analysis of the skin, and Dr. Benjamin D. Yu (University of California, San Diego) for the gift of the A388 cell line.

## Author contribution

Conceptualization, K.B. and C.J.; Methodology, K.B. and C.J.; Investigation, K.B., B.D., S.K., R.K.Z., R.S., T.M., R.D. and A.G.; Validation, K.B., B.D. and S.K.; Formal Analysis, R.D. and J.S., P.K.; Resources, C.J., S.K, R.S., A.G., and Y.W.H., Writing – Original Draft, K.B. and C.J.; Funding Acquisition, C.J. and S.K.; Supervision, C.J. and S.K.

