## Supplementary Information for "Snail maintains the stem/progenitor state of skin epithelial cells and carcinomas through the autocrine effect of the matricellular protein Mindin"

### Figure S1

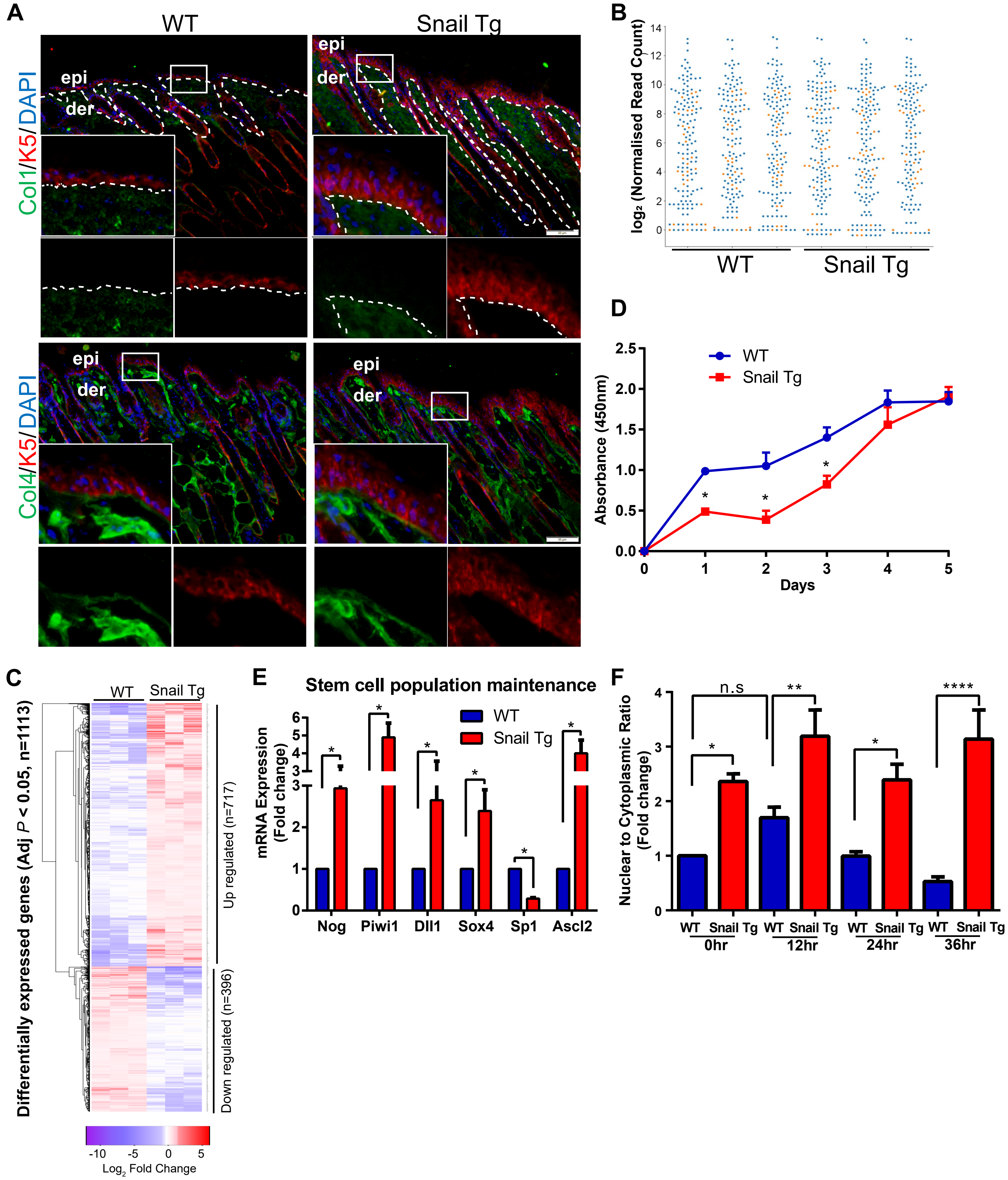


**Figure S1: Overexpression of Snail reinforces the stem/progenitor characteristics in vitro. (Related to Figure 1)**

(A) Immunofluorescence assay representing intact compartmentalization of cells in the skin. Top row: Collagen 1 (green) to mark the mesenchymal cells, Keratin 5 (red) to mark the basal epidermal keratinocytes and outer root sheath of hair follicles in WT and Snail Tg skin. Bottom row: Collagen 4 (green) marks the mesenchymal cells and keratin 5 (red) marks the basal epidermal keratinocytes. Insets in each panel are magnified views of the boxed areas of the epidermis. Scale bar: 50µm
(B) The gene expression values of the EMT signature genes in the WT and Snail Tg keratinocytes.
(C) Heat map of the differentially expressed genes in Snail Tg keratinocytes compared to its WT counterpart.
(D) Graph representing the proliferative capacity of WT and Snail Tg keratinocytes (n=3).
(E) qPCR of genes involved in stem cell population maintenance in Snail Tg cells compared to WT counterpart (n=3).
(F) Nuclear to cytoplasmic ratio of differentiating WT and Snail Tg keratinocytes in the calcium switch assay.
Error bars depict ± SEM. * $P < 0.05$, ** $P < 0.01$, ****$P < 0.0001$ and n.s=not significant.

### Figure S2

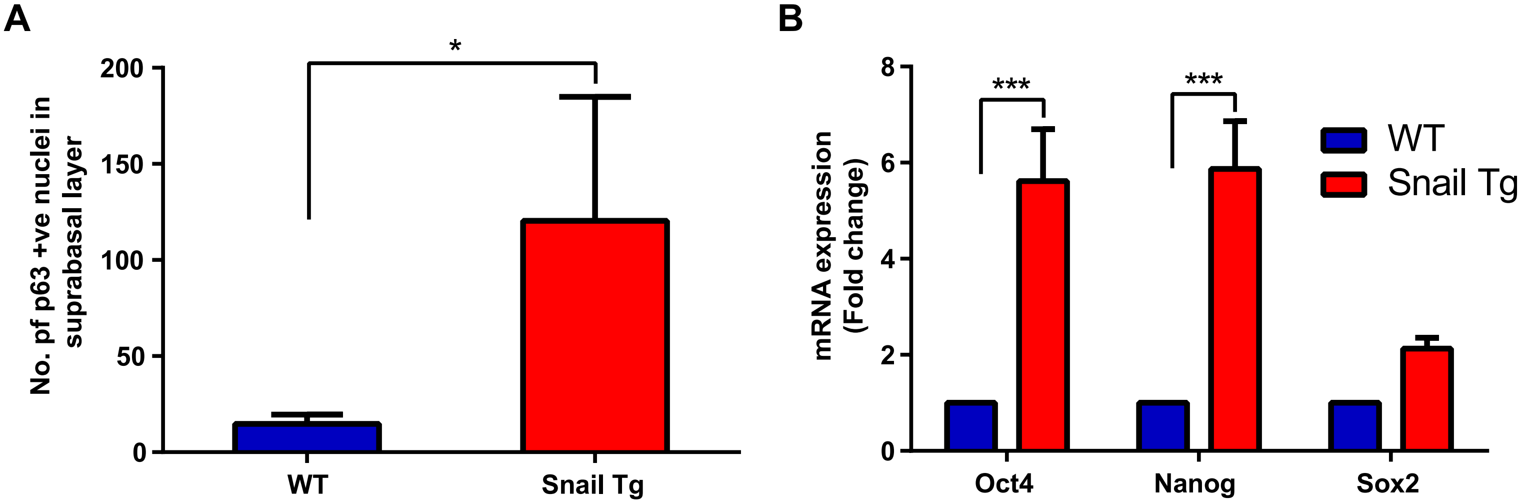


**Figure S2: Epidermal Snail expression expands the epidermal stem/progenitor cell population in vivo (Related to Figure 2)**

(A) Quantification of p63 positive nuclei in the suprabasal layers of the WT and Snail Tg epidermis (n=3).
(B) mRNA expression of stem cell genes Oct4, Sox2 and Nanog in WT and Snail Tg skin quantified by qPCR (n=3).
Error bars depict ± SEM. * $P < 0.05$ and *** $P < 0.001$

### Figure S3

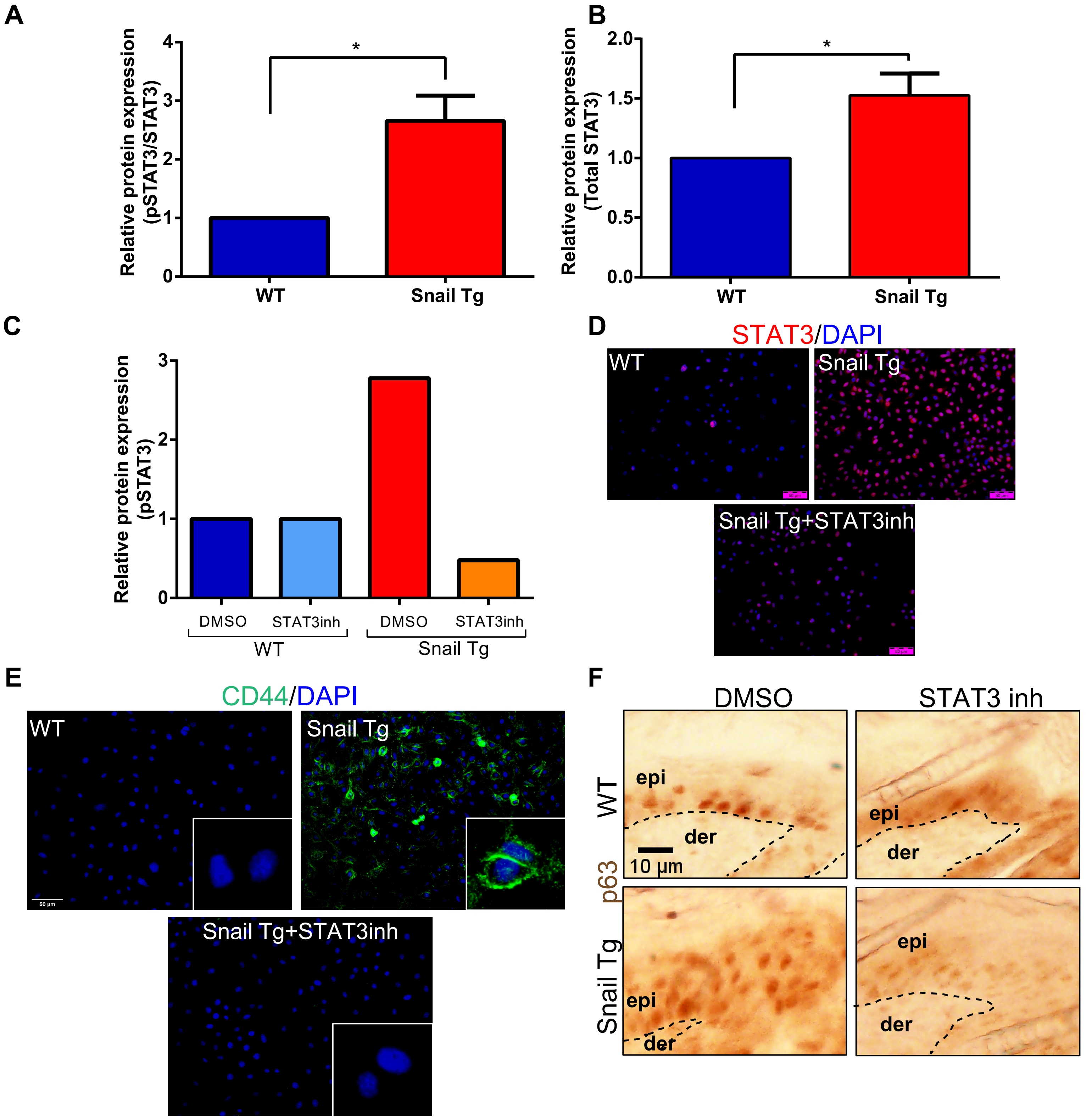


**Figure S3: Snail mediated stem/progenitor cell maintenance is dependent upon STAT3 activation (Related to Figure 3)**

(A) Quantification of the western blot analysis for pSTAT3 to STAT3 levels in WT and Snail Tg keratinocytes (n=3).
(B) Quantification of total STAT3 levels in WT and Snail Tg keratinocytes (n=3).
(C) pSTAT3 levels quantified by western blot analysis in WT and Snail Tg keratinocytes treated with STAT3 inhibitor, BP-1-102.
(D) Immunofluorescence for STAT3 in red and DAPI in blue in WT, Snail Tg and Snail Tg cells treated with STAT3 inhibitor. Scale bar: 50μm
(E) Immunofluorescence of stem cell marker, CD44 in green in WT, Snail Tg and Snail Tg +STAT3 inhibited keratinocytes. Scale bar: 50μm
(F) Immunohistochemistry of p63 in WT and Snail Tg skin treated with either DMSO or STAT3 inhibitor. Scale bar: 10μm
Error bars depict ± SEM. $^*P < 0.05$

### Figure S4

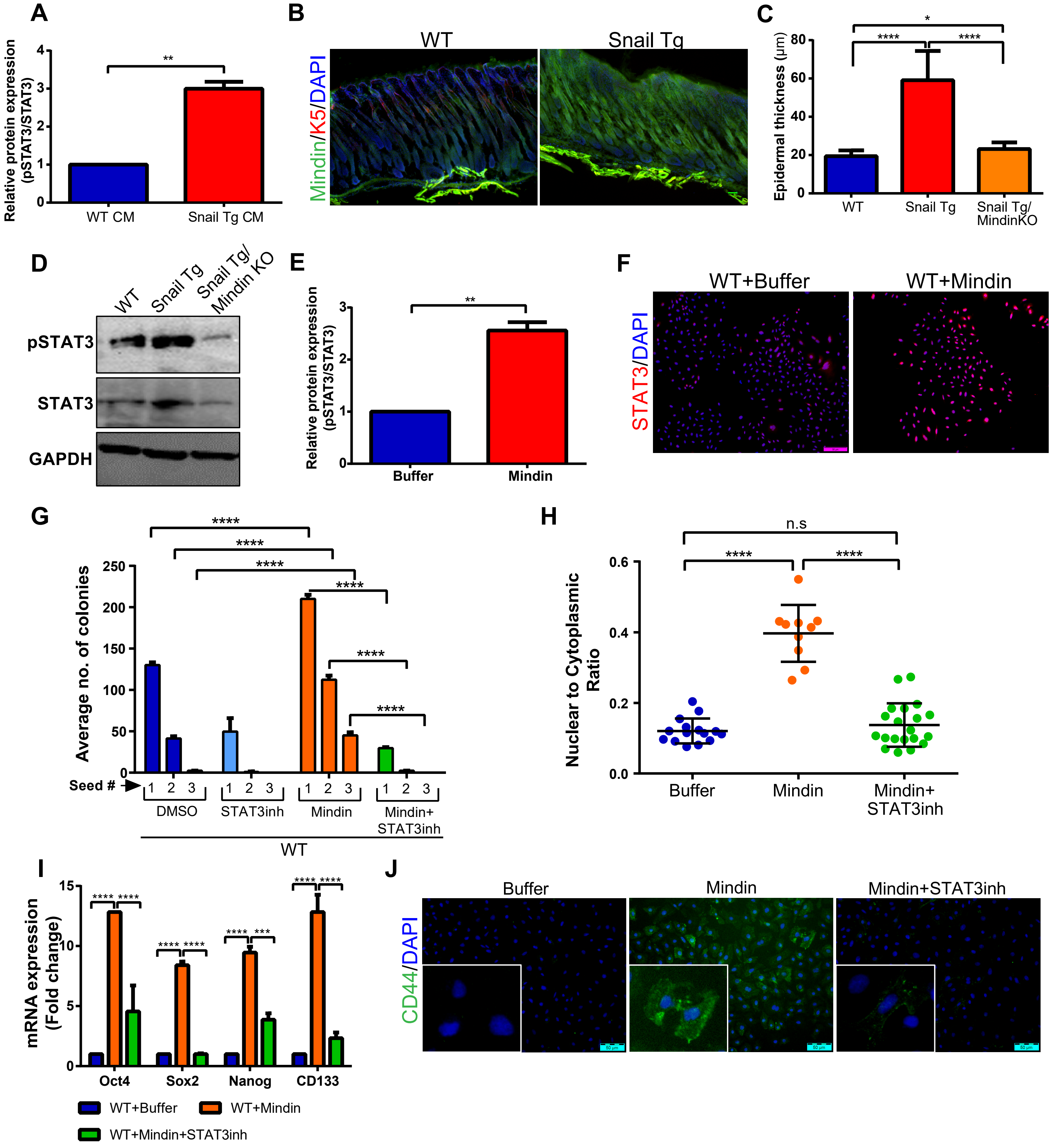


**Figure S4: Secreted Mindin by Snail Tg keratinocytes activates STAT3 in an autocrine fashion (Related to Figure 4)**

(A) Western blot quantification of pSTAT3/STAT3 levels in keratinocytes treated with conditioned media (CM) either from WT or Snail Tg keratinocytes (n=3).
(B) WT and Snail Tg skin analysed for secreted Mindin (green). Absence of keratin 5 (red) staining was used as a positive control for non-permeabilization. Scale bar: 40μm.
(C) Epidermal thickness quantified on WT, Snail Tg and Snail Tg/Mindin KO skin.
(D) Western blot analysis of pSTAT3 and total STAT3 in WT, Snail Tg, and Snail Tg/Mindin KO keratinocytes. GAPDH was used as loading control (n=3).
(E) Western blot quantification of pSTAT3/STAT3 levels in keratinocytes treated with recombinant Mindin protein along with its buffer control (n=3).
(F) Immunofluorescence of STAT3 in red on keratinocytes treated with either recombinant Mindin protein or buffer. Scale bar: 50μm.
(G) Average number of colonies formed by wild type keratinocytes treated with either Mindin or Mindin and STAT3 inhibitor along with its respective controls (n=3).
(H) Quantification of the nuclear to cytoplasmic ratio of WT keratinocytes treated with either Mindin or both Mindin and STAT3 inhibitor along with it's buffer control.
(I) mRNA expression of stem cell genes Oct4, Sox2, Nanog, CD133 in WT keratinocytes treated with either Mindin or both Mindin and STAT3 inhibitor (n=3).
(J) Immunofluorescence of CD44 in green on WT keratinocytes treated with either Mindin or both Mindin and STAT3 inhibitor. Scale bar: 50μm.
Error bars depict ± SEM. * $P < 0.05$, ** $P < 0.01$, *** $P < 0.001$, ****$P < 0.0001$ and n.s=not significant.

### Figure S5

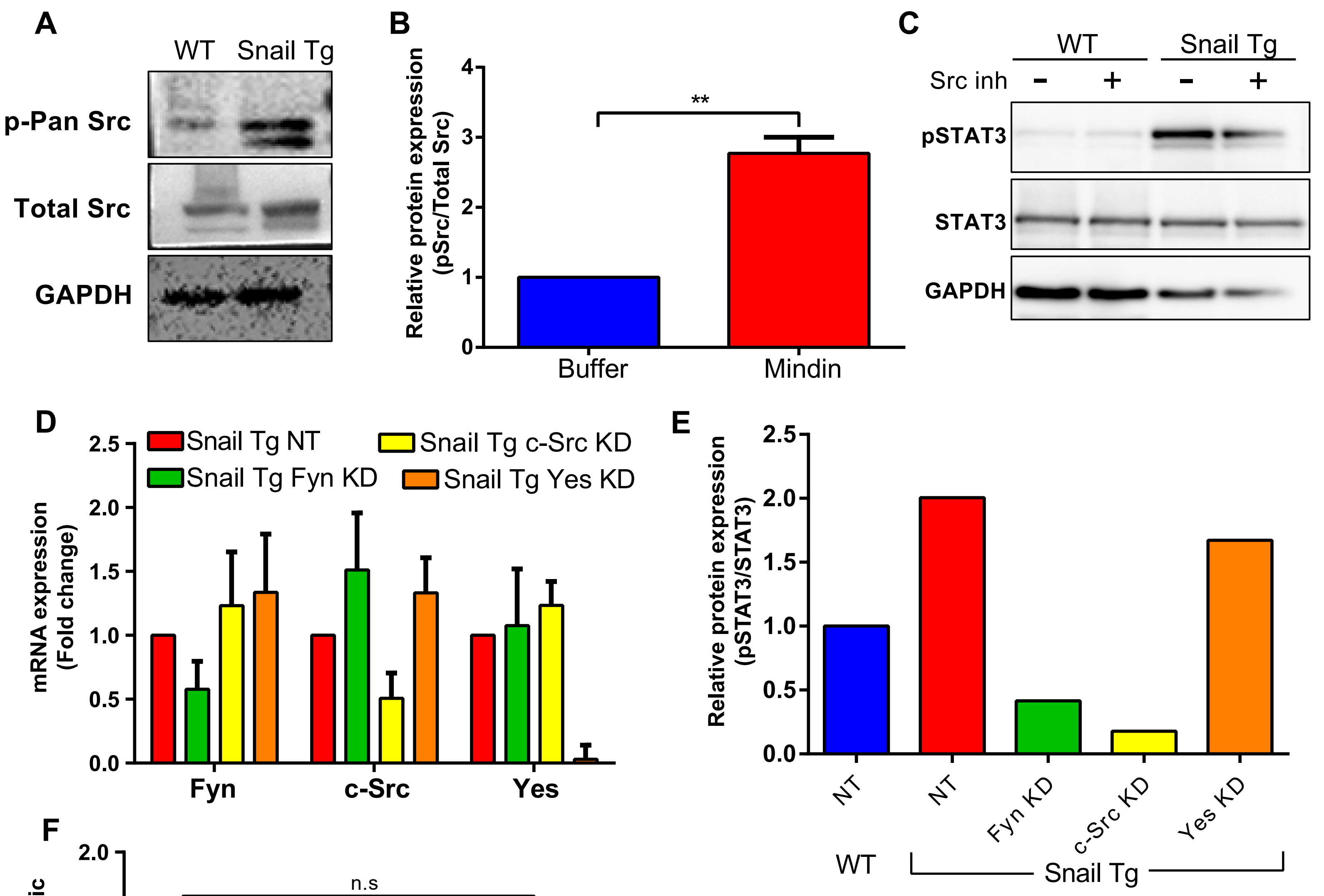


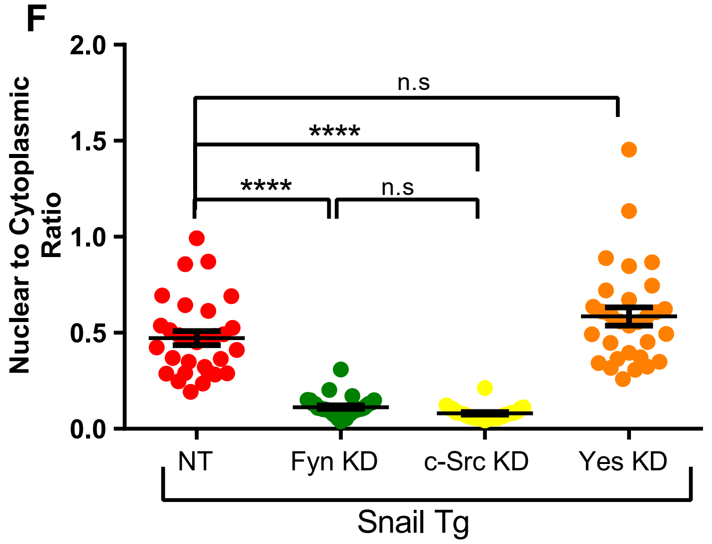


**G**

| SFK Kinase | Colony formation assay | Stem cell gene expression | NCR | STAT3 activation |
|---|---|---|---|---|
| Fyn | + | + | + | + |
| c-Src | + | - | + | + |
| Yes | +/- | + | - | - |

**Figure S5. Src Family of Kinases mediate the Mindin dependent activation of STAT3 (Related to Figure 5)**

(A) Western blot for phospho pan Src and total Src in WT and Snail Tg keratinocytes (n=3). GAPDH was used as the loading control.
(B) Quantification of western blot analysis of phospho pan Src to total Src in WT keratinocytes treated with Mindin and its buffer as control (n=3).
(C) Western blot for pSTAT3 and total STAT3 in WT and Snail Tg keratinocytes treated with Src inhibitor and its buffer control. GAPDH was used as the loading control.
(D) qPCR validation to show the specificity of the shRNA mediated Fyn, c-Src and Yes knock downs with NT as control (n=3).
(E) Quantification of western blot analysis of phospho STAT3 to total STAT3 in the SFK KDs with Snail Tg NT compared to WT as control.
(F) Quantification of the nuclear to cytoplasmic ratio of the SFK KDs relative to the Snail Tg NT as control.
(G) Table summarizing the contribution of each of the three SFKs in the Snail mediated stemness associated with keratinocytes.
Error bars depict ± SEM. ** $P < 0.01$, ****$P < 0.0001$ and n.s=not significant.

### Figure S6

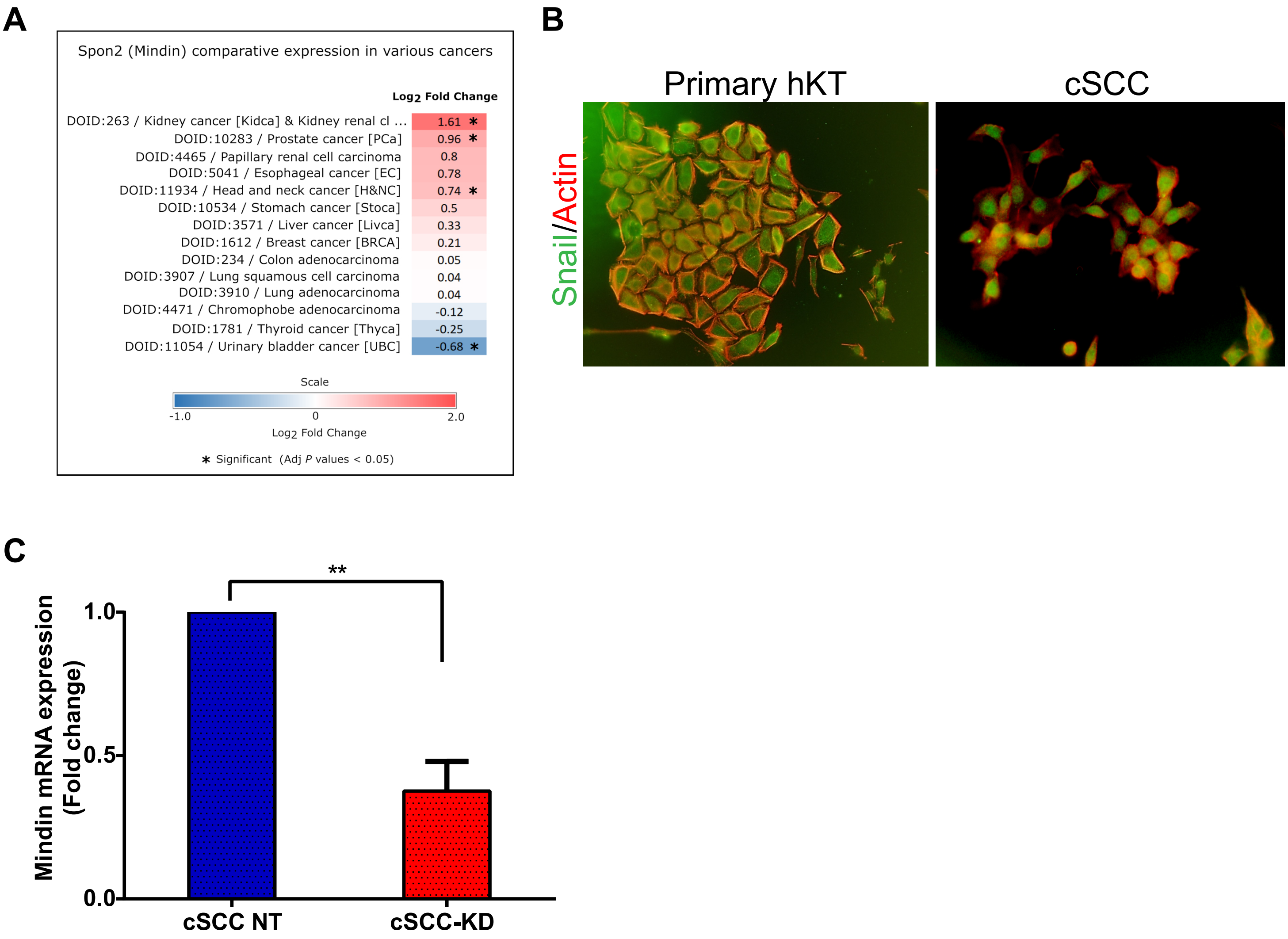


**Figure S6. Mindin is required for the maintenance of stem/progenitor characteristics in cutaneous squamous cell carcinoma cells (Related to Figure 6)**

(A) Expression pattern of Mindin in various carcinomas.
(B) Immunofluorescence for Snail in green and actin in red shows nuclear localization of Snail in cSCC cells compared to primary human keratinocytes.
(C) qPCR validation of CRISPR-Cas9 mediated Mindin knock down (KD) in cSCC cells. No Target (NT) was used as a control (n=3). Error bars depict ± SEM. ** P < 0.01

### Figure S7

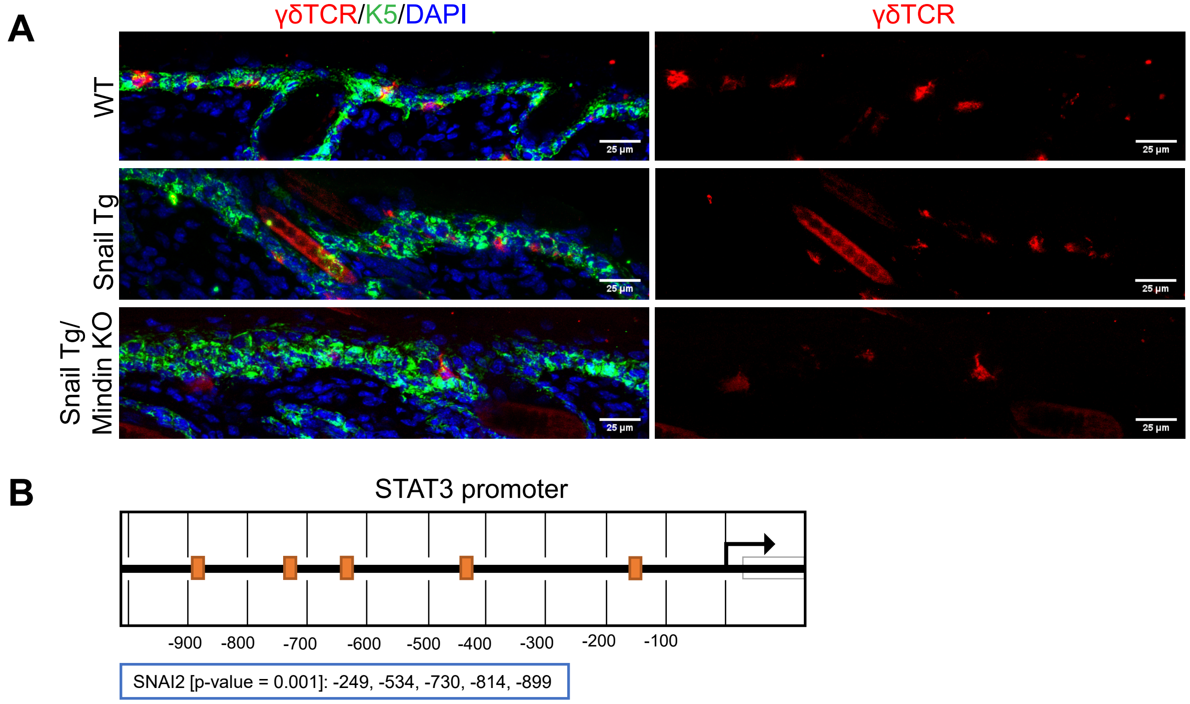


**Figure S7:**

(A) Immunofluorescence of γδTCR (red) to mark the γδT-cells, K5 (green) that marks the basal layer of the epidermis in WT, Snail Tg and Snail Tg/Mindin KO skin. Scale bar: 25µm.
(B) E-box sequence for Snail binding on the STAT3 promoter region shown in orange.

**Supplementary Table 1**

| | Mouse Gene | Forward primer sequence | Reverse primer sequence |
|---|---|---|---|
| 1 | Noggin | GCCAGCACTATCTACACATCC | GCGTCTCGTTCAGATCCTTCTC |
| 2 | Piwil2 | TTGGCCTCAAGCTCCTAGAC | GAACATGGACACCAAACCTACA |
| 3 | Dll1 | CAGGACCTTCTTTCGCGTATG | AAGGGGAATCGGATGGGGTT |
| 4 | Sox4 | GACAGCGACAAGATTCCGTTC | GTTGCCCGACTTCACCTTC |
| 5 | Spi1 | AGAAGCTGATGGCTTGGAGC | GCGAATCTTTTTCTTGCTGCC |
| 6 | Ascl2 | CCGTGAAGGTGCAAACGTC | CCCTGCTACGAGTTCTGGTG |
| 7 | ZFP703 | GCTGGATCTAACCCAAGGACA | GACAGCGGTTGCAGGTACTC |
| 8 | IGIF1R | GTGGGGGCTCGTGTTTCTC | GATCACCGTGCAGTTTTCCA |
| 9 | TNC | GCAGTGAAAAGCGGTGTCC | CTTCTCCGGTATAGCCCTCGT |
| 10 | Prrx1 | AGCAGACGAAAGTGTGGGC | TCAGAGTTCAACTGGTCATTGTC |
| 11 | PTGS2 | TTCCAATCCATGTCAAAACCGT | AGTCCGGGTACAGTCACACTT |
| 12 | IGFBP3 | CACACCGAGTGACCGATTCC | GTGTCTGTGCTTTGAGACTCAT |
| 13 | PTPRK | TGGATTCACTGGTCGTGATTG | TCGCAAGGATAACTCAGGACTT |
| 14 | PTEN | TGGATTCGACTTAGACTTGACCT | GCGGTGTCATAATGTCTCTCAG |
| 15 | PRKX | GCGACCGTAAAAGATCCAGAC | GTTCCTGTTTCAGGCGGATGA |
| 16 | Keratin 5 | GCCCACAGAGACTGCTTCTT | TTGGGATTCGTTCCCTTCCG |
| 17 | Involucrin | ATGTCCCATCAACACACACTG | TGGAGTTGGTTGCTTTGCTTG |
| 18 | Loricrin | AGGCAGTCTTGAAGAATCCAGA | GCCAGCTTTAGCACCAGTAGTAA |
| 19 | Pai 1 | GCTGCACCCTTTGAGAAAGA | GCCAGGGTTGCACTAAACAT |
| 20 | Oct4 | CACCATCTGTCGCTTCGAGG | AGGGTCTCCGATTTGCATATCT |
| 21 | Sox2 | GGCAGCTACAGCATGATGCAGGAGC | CTGGTCATGGAGTTGTACTGCAGG |
| 22 | Nanog | AGGGTCTGCTACTGAGATGCTCTG | CAACCACTGGTTTTTCTGCCACCG |
| 23 | CD133 | TCCGGTGTCATAGCTGGGTA | CACTAGGTCACAACCACCCC |
| 24 | Mindin | ATGGAAAACGTGAGTCTTGCC | TGATGCTGTATCTAGCCAGAGG |
| 25 | PITX1 | ATCGTCCGACGCTGATCTG | GCTTGTGAAGTGAGTGCGTT |
| 26 | GAPDH | AGGTCGGTGTGAACGGATTTG | TGTAGACCATGTAGTTGAGGTCA |
| | **Human Gene** | **Forward primer sequence** | **Reverse primer sequence** |
| 1 | Mindin | CGCTGGACCTGTACCCCTA | AGGAGGACGTTATCTCGGTCA |
| 2 | Oct4 | CTTGAATCCCGAATGGAAAGGG | GTGTATATCCCAGGGTGATCCTC |
| 3 | Sox2 | GCCGAGTGGAAACTTTTGTCG | GGCAGCGTGTACTTATCCTTCT |
| 4 | Nanog | TTTGTGGGCCTGAAGAAAACT | AGGGCTGTCCTGAATAAGCAG |
| 5 | CD133 | AGTCGGAAACTGGCAGATAGC | GGTAGTGTTGTACTGGGCCAAT |
| 6 | PITX1 | GTTCAGCGGCCTAGTGCAG | CGGGCTCATGGAGTTGAAGAA |
| 7 | Nrf2 | TCAGCGACGGAAAGAGTATGA | CCACTGGTTTCTGACTGGATGT |
| 8 | Klf4 | CCCACATGAAGCGACTTCCC | CAGGTCCAGGAGATCGTTGAA |
| 9 | Actin | TCCTTCCTGGGCATGGAGT | AGCACTGTGTTGGCGTACAG |

**Supplementary Table 2**

| Designation | Source or reference | Identifiers | Additional information |
|---|---|---|---|
| anti- total STAT3 | CST | 4904 | 1:1000 (WB), 1:200 (IF) |
| anti-phospho STAT3 | CST | 9145S | 1:1000 (WB), 1:200 (IF) |
| anti-p63 | Abcam | ab97865 | 1:1000 (WB), 1:200 (IF) |
| anti-GAPDH | CST | 5174S | 1:2000 (WB) |
| anti-Tubulin | Sigma | T5326-200UL | 1:5000 (WB), 1:200 (IF) |
| WGA | Invitrogen | W32466 | 1:200 (IF) |
| DAPI | Molecular Probes | | 1:1000 (IF) |
| anti-Keratin 5 | Jamora lab generated | | 1:200 (IF) |
| anti-CD49f | BD | 561894 | 1:100 (IF) |
| anti-CD44 | BD | 550538 | 1:200 (IF) |
| anti-Mindin | Santa Cruz | SC49050 | 1:75 (IF) |
| anti-total Src | CST | 2123 | 1:1000 (WB) |
| anti-phospho Src | CST | 6943 | 1:1000 (WB) |
| Alexa 488- or 555-secondaries | Molecular Probes | | (1:400) |
| STAT3 inhibitor | Millipore | 573132-10MG | |
| Src inhibitor | Millipore | 567805 | |
| Matrigel | BD Biosciences | 354277 | |
| GraphPad | Prism | | ver.6 for statistical analysis |

**Supplementary Table 3**

| shRNA Construct | Sequence |
|---|---|
| ULTRA-3238565 Src | TGCTGTTGACAGTGAGCGCTGTGACAGAGTACATGAACAATAGTGAAGCCACAG ATGTATTGTTCATGTACTCTGTCACAATGCCTACTGCCTCGGA 1 |
| ULTRA-3238564 Src | TGCTGTTGACAGTGAGCGCCCACGAGGGTTGCCATCAAAATAGTGAAGCCACAG ATGTATTTTGATGGCAACCCTCGTGGTTGCCTACTGCCTCGGA 1 |
| ULTRA-3238562 Src | TGCTGTTGACAGTGAGCGAAAGAACCCATTTACATTGTGATAGTGAAGCCACAGA TGTATCACAATGTAAATGGGTTCTTCTGCCTACTGCCTCGGA 1 |
| ULTRA-3216477 Fyn | TGCTGTTGACAGTGAGCGAAAGGTTCACAATCAAGTCTGATAGTGAAGCCACAG ATGTATCAGACTTGATTGTGAACCTTCTGCCTACTGCCTCGGA 1 |
| ULTRA-3216480 Fyn | TGCTGTTGACAGTGAGCGCCTGCGATCAGCAAACATTCTATAGTGAAGCCACAG ATGTATAGAATGTTTGCTGATCGCAGATGCCTACTGCCTCGGA 1 |
| ULTRA-3216478 Fyn | TGCTGTTGACAGTGAGCGAGGACCACGTCAAACATTATAATAGTGAAGCCACAG ATGTATTATAATGTTTGACGTGGTCCCTGCCTACTGCCTCGGA 1 |
| ULTRA-3247308 Yes1 | TGCTGTTGACAGTGAGCGCAAGAAGGAGATGGAAAGTATATAGTGAAGCCACAG ATGTATATACTTTCCATCTCCTTCTTTTGCCTACTGCCTCGGA 1 |
| ULTRA-3247307 Yes1 | TGCTGTTGACAGTGAGCGAAAGGGTGAACGATTTCAAATATAGTGAAGCCACAG ATGTATATTTGAAATCGTTCACCCTTCTGCCTACTGCCTCGGA 1 |
| ULTRA-3247309 Yes1 | TGCTGTTGACAGTGAGCGCCCTGATGAAAGACCAACATTATAGTGAAGCCACAG ATGTATAATGTTGGTCTTTCATCAGGATGCCTACTGCCTCGGA 1 |
| TLNSU1440 Non-targeting control #1 | TGCTGTTGACAGTGAGCGCCCGGCTGAAGAGCCTGATCAATAGTGAAGCCACAG ATGTATTGATCAGGCTCTTCAGCCGGTTGCCTACTGCCTCGGA 5 |
